# The two key substitutions in the chromophore environment of mKate2 to produce an enhanced FusionRed-like red fluorescent protein

**DOI:** 10.1101/2023.12.05.570295

**Authors:** Dmitry A. Ruchkin, Alexey S. Gavrikov, Danila V. Kolesov, Andrey Yu. Gorokhovatsky, Tatiana V. Chepurnykh, Alexander S. Mishin, Eugene G. Maksimov, Nadya V. Pletneva, Vladimir Z. Pletnev, Anastasiia M. Pavlova, Vladimir A. Nikitin, Konstantin A. Lukyanov, Alexey M. Bogdanov

**Affiliations:** Shemyakin-Ovchinnikov Institute of Bioorganic Chemistry, Moscow 117997, Russia; Faculty of Biology, M.V. Lomonosov Moscow State University, 119992 Moscow, Russia; Pirogov Russian National Research Medical University, 117997 Moscow, Russia; Department of Photonics, İzmir Institute of Technology, 35430 İzmir, Turkey

## Abstract

Red fluorescent proteins (RFPs) are often probes of choice for living tissue microscopy and whole-body imaging. When choosing a specific RFP variant, the priority may be focused on fluorescence brightness, maturation rate, monomericity, excitation/emission wavelengths, and low toxicity, which are rarely combined in optimal way in a single protein. If the additional requirements such as prolonged fluorescence lifetime and/or blinking ability are applied, the available probe’s repertoire could become surprisingly narrow. Since the whole diversity of the conventional single-component RFPs belongs to just a few phylogenetic lines (with DsRed-, eqFP578- and eqFP611-derived being the major ones), it is not unexpected that their advantageous properties are split between close homologs. In such cases, a systematic mutagenetic analysis focused on variant-specific amino acid residues can shed light on the origins of sibling RFPs’ distinctness and might be beneficial for consolidation of their strengths in new RFP variants. For instance, the protein FusionRed, although being efficient in the fluorescence labeling due its good monomericity and low cytotoxicity, has undergone a considerable loss in fluorescence brightness/lifetime compared to the parental mKate2. In this contribution, we describe a fast-maturing monomeric RFP designed semi-rationally based on the mKate2 and FusionRed templates, outperforming both its parents in molecular brightness, having extended fluorescence lifetime, and showing spontaneous blinking pattern promising for nanoscopy use.

## Introduction

Current bioimaging techniques recruit a vast diversity of fluorescent probes, among which genetically encoded fluorophores, such as the fluorescent proteins (FPs), are in favor, enabling highly specific intracellular labeling, live-cell superresolution, and fluorescence lifetime imaging microscopy (FLIM), etc. [1, 2]. In turn, red fluorescent proteins (RFPs), a polyphyletic group [3–6] of anthozoan FPs emitting in the red part of the spectrum, are of a special relevance for the whole-body and/or deep-tissue imaging owing to their enhanced detectability within a ‘window of optical transparency’ characterizing a local absorption minimum of the animal tissues at the wavelengths of ∼600-1200 nm [7–9]. Among existing RFP variants, FusionRed [10] is often a preferred probe for live-cell imaging (including visualization of fine subcellular structures) thanks to its ‘supermonomericity’, i.e., an ability to maintain highly monomeric state even at the great local concentrations typical for the specialized localizations within the mammalian cells [11, 12], low acid sensitivity and toxicity. It is thus supposed to be used as a probe fused to the proteins of interest without affecting their natural activities and spatial structures, or functioning as a fluorescent core of the genetically encoded indicators [13–16]. In the meantime, there are definite drawbacks which take from the value of FusionRed as a multipurpose fluorescence tag and call for further improvement of this RFP. Thus, an issue of the modest molecular brightness possessed by FusionRed has been extensively addressed in the elegant studies by the Jimenez lab, where both a directed evolution [17] and semi-rational design [18] were utilized to engineer the brighter variants of FusionRed (specifically, the FusionRed-MQV [19] shows ∼4-fold molecular brightness advantage over the parental RFP, though its emission peak has a 20-nm hypsochromic shift). The high-resolution spatial structure of FusionRed revealed that almost a half of its molecules carries an immature chromophore; this feature reduces effective brightness (well under the level expected based on the measured molecular brightness) of FusionRed as a fluorescence probe, providing further space for improvement via structure-based design of daughter RFP variants.

Importantly, FusionRed is a descendant of the mKate2 protein [20] emitting at 633 nm (its emission maximum is 25-nm red-shifted relative to that of FusionRed), and is currently the brightest monomeric far-red FP. FusionRed differs from mKate2 in 17 amino acid substitutions introduced semi-rationally, through the several consecutive rounds of mutagenesis [10]. Hence, there is no unified picture describing a particular role of every substitution, specifically, structural determination of the spectral differences (including extinction coefficient, fluorescence quantum yield and lifetime, excitation/emission maxima positions) between the FusionRed and mKate2 proteins is not clear enough. Based on the analysis of the spatial structure of FusionRed [21], one can consider an essential role of 3 residues from the chromophore environment, Arg/Lys-67, Cys/Ala-158, His/Arg-197 (FusionRed/mKate2, respectively). Here, we study an influence of these residues on the properties of both proteins systematically by an exhaustive reciprocal site-directed mutagenesis. Among representatives of the library obtained, there is a remarkable variant – mKate2-K67R/R197H – which shows a striking similarity in its steady-state absorption and fluorescence spectra to those of FusionRed and is 2.2-fold brighter than the latter. This RFP inherits the advantages of both sister proteins, namely, it demonstrates a monophasic fluorescence decay like mKate2, and shows good performance as a fusion tag like FusionRed. Interestingly, a purified mKate2-K67R/R197H possesses a well-marked pattern of spontaneous fluorescence blinking that might be promising for superresolution microscopy use.

## Materials and methods

### Site-directed mutagenesis

To obtain the site-specific mutants of mKate2 and FusionRed, a modified IVA-cloning [22] procedure was applied. The genes of taken RFPs cloned in the pQE-30 vector backbone (Qiagen, Germantown, Maryland, USA) using BamHI/HindIII endonuclease sites were used as primary templates. The forward oligonucleotides were designed to have a 5’-terminal 15-20 nt length region with homology to the template DNA (needed to provide bacterial recombination), followed by a triplet with a mutation of interest and a 3’-terminal priming region designed to anneal at a temperature of 60°C. The reverse oligonucleotides consisted of a similar recombination-guide part of 15-20 nt and a 3’-terminal priming sequence; both made up annealing temperature of 60°C when possible. In cases of higher calculated annealing temperatures, the 5’-end fragment was considered partially annealing. The 3’ and 5’ terminal bases of both primers were selected not to pair complementarily in order to avoid self-annealing of long oligonucleotides if possible; simultaneously, the terminal 3’-nucleotides of both primers were preferably selected to form strong complementary pair with the template sequence. The reverse primer could never anneal to the forward with 3’-terminus resulting in a blunt-end. The PCR was carried out using a standard Phusion Polymerase (ThermoFisher, Waltham, Massachusetts, USA) protocol and lasted 35 cycles; the template DNA made up for a total of 50 ng per reaction. The primers used had the following sequences:

a. FusionRed-R67K: Forward - 5’- agcttcatgtacggcagcaaaaccttcatcaagcaccct ccgg-3’ Reverse - 5’-gctgccgtacatgaagctggtag-3’
b. FusionRed-C158A: The mutant was engineered in the previous study [21] Forward - 5’-cggcggcctggaaggcgcagcagacatggccctgaa gctcg-3’ Reverse - 5’-tgcgccttccaggccgccgtcagcggggtacatcgtctc g-3’
c. FusionRed-H197R: Forward - 5’-ggcgtctacaacgtggacagaagactggaaagaatcaaggaggc-3’ Reverse - 5’-gtccacgttgtagacgccgggcatcttgaggttcgtagc g-3’
d. mKate2-K67R: Forward - 5’-agcttcatgtacggcagcagaaccttcatcaaccacac ccaggg-3’ Reverse - 5’-tgctgccgtacatgaagctggtag-3’
e. mKate2-A158C: The mutant was engineered in the previous study [21] Forward - 5’-ggcctggaaggcagatgcgacatggccctgaagctcg -3’ Reverse - tctgccttccaggccgccgtcagcggggtacag-3’
f. Kate2-R197H: 5’- Forward - 5’-ggcgtctactatgtggaccacagactggaaagaatcaa ggaggc-3’ Reverse - 5’-gtccacatagtagacgccgggcatcttgaggttcttagc g-3’

The PCR products were reprecipitated and treated with DpnI restriction endonuclease to remove the initial template DNA. For transformation (needed for constructs assembly), 700 ng of PCR product was taken per one 100µl aliquot of E.coli XL1-Blue competent cells (Evrogen, Moscow, Russian Federation).

### Protein expression and purification

The FP variants were expressed in E.coli XL1-Blue strain for 72 hours at 37ºC. After centrifugation, bacterial biomass was resuspended in PBS (GIBCO, ThermoFisher Scientific, Waltham, Massachusetts, USA) pH 7.4 and treated with ultrasound by Sonics Dismembrator (Fisher Scientific, Pittsburgh, Pennsylvania, USA). The proteins were then purified using TALON metal-affinity resin (Clontech, Mountain View, California, USA) previously added and washed in PBS according to the manufacturer’s protocol, and solubilized using 0.3 mM imidazole (pH 8.0). Protein eluates were then desalted and concentrated by ultrafiltration with Amicon Ultra 0.5 10K (Merck Millipore, Burlington, Massachusetts, USA) columns. The obtained concentrated protein solution (typically ∼5 mg/ml) was ready to be used for SDS-PAGE analysis or spectroscopy or shortly stored at 4ºC until used.

### Steady-state absorption and fluorescence spectroscopy

The absorbance and fluorescence spectra were recorded using a Cary100 UV/VIS spectrophotometer and a Cary Eclipse fluorescence spectrophotometer (Agilent Technologies, Santa Clara, California, USA), respectively. In all cases, a protein solution in PBS (pH 7.4) was used. The fluorescence quantum yields and extinction coefficients were determined as described earlier [21].

### Monomericity testing

#### Gel-filtration

Gel-filtration experiments were performed using a Superdex® 200 Increase 10/300 GL column (Cytiva, Uppsala, Sweden) equilibrated with 20 mM sodium phosphate buffer (pH 7.4) containing 150 mM NaCl at 24 °C and at a flow rate of 0.75 ml/min. The column was connected to an Agilent 1260 Bio-Inert LC system equipped with in-line Agilent 1260 diode array detector and Agilent 1260 fluorescence detector and calibrated using cytochrome C (12.4 kDa), carbonic anhydrase (29 kDa), bovine serum albumin (66 kDa), alcohol dehydrogenase (150 kDa), b-amilase (200 kDa) and ferritin (450 kDa). For the calibration details, see Fig. S6 and table S1. The equipment was controlled by Agilent OpenLAB CDS ChemStation Edition C.01.07 SR3 software.

#### OSER assay

The OSER assay was carried out in two variants. The first one, based on HeLa cells, was similar to that described in [12]. The cells were transfected with FuGENE® HD Transfection Reagent (Promega, Woods Hollow Road, Madison, USA) following the commercial protocol. Images were acquired with wide-field fluorescence microscopy using a modified Leica 6000LX inverted microscope equipped with an mCherry filter cube (see Widefield fluorescence microscopy section). Processing of images was performed using the Fiji ImageJ distribution (version 2.9.0/1.54b). Whorl-like structures were then detected according to the guidance by Constantini et al. [23]. Due to the lack of whorl-like structures in more than 80% of transfected HeLa cells and the resulting high difficulty of capturing enough whorl-possessing HeLa cells for valid statistical analysis, the calculation of the mean fluorescent intensities of nuclear envelope to whorl-like structures ratios was not carried out.

The second variant of the OSER assay was similar to that described in [23]. U2OS cells were transfected with PEI (Sigma-Aldrich, Saint Louis, Missouri, USA). Observation took place 18 hours after transfection; images were analyzed using the same Fiji ImageJ software to obtain the mean NE and OSER signals. Not less than three linear ROIs for NE were traced for each cell using the ‘straight’ tracing instrument; for OSER ROIs, the ‘freehand’ instrument was used. Ratios were calculated using GraphPad Prism10.

### Engineering of mammalian constructs

Mammalian expression plasmids encoding fusions of Diogenes with vimentin (Vimentin-FR2), lifeact (lifeact-FR2), ensconsin (ensconsin-FR2) and cytokeratin (FR2-cytokeratin) as well as the fusion with cytoplasmic end of an endoplasmic reticulum signal anchor membrane protein (CytERM; used in the OSER assay) were assembled using Golden Gate cloning following MoClo standard procedure [24–26]. Each transcriptional unit for mammalian expression included CMV promoter, coding sequence for the fusion protein, and the SV40 terminator. All Golden Gate cloning reactions were performed in the T4 ligase buffer (SibEnzyme, Moscow, Russia) with 10 U of T4 ligase, 20 U of either BsaI or BpiI restriction endonucleases (ThermoFisher, Waltham, Massachusetts, USA), and 100-200 ng of DNA of each DNA fragment. The assembly reactions were performed under the following conditions: 30 cycles of 37°C and 16°C incubations (90 sec at 37°C, 180 sec at 16°C).

### Widefield fluorescence microscopy

Widefield fluorescence microscopy was performed with a Leica 6000LX inverted microscope, equipped with a Leica HCX PL APO 100X/1.40–0.70NA oil immersion objective, Zyla sCMOS camera (Andor, Oxford, UK), and CoolLED pE-300 light source. An mCherry cube filter set (Leica, Wetzlar, Germany) was used (excitation filter: 560/40, emission filter: 630/75). Typical illumination power ranged from 1 to 5 W/cm^2^ with exposure times ranging from 50 to 150 ms.

### pH-stability measurement

The range of premade buffer solutions with pH from 3 to 10.55 was used to prepare the protein samples; the solutions contained 130 mM KCl, 30 mM NaCl, 0.5 mM MgCl_2_, 0.2 mM EGTA and 30 mM HCl– NaH_2_C_6_H_5_O_7_ (pH 3.0–4.5) or 15 mM KH_2_PO_4_–Na_2_HPO_4_ (pH 5.0–7.5) or 20 mM Na_2_B_4_O_7_–HCl/NaOH (pH 8.0–11.0) [27]. Each probe contained 5 μg/mL of the purified and desalted RFP. For each sample, the emission spectra were measured with a Cary Eclipse Fluorescence Spectrometer twice for each of three temporary points (immediately after preparation, in 3 min and in 5 min) with a total of 6 measurements per sample in the spectral range from 560 nm to 700 nm at λ_ex_ = 540 nm using 5 nm ex/em slit, equal photomultiplier (PMT) voltage and scanning speed values. Fluorescence intensity values at emission maxima were averaged from 6 reads. The averaged data from all pH points for each RFP were normalized to the maximum value within the set and plotted on a graph with standard deviations. The sigmoidal regions of the graphs were fitted (4PL logistic curve, 95% confidence, n=6) in GraphPad Prism10; pKa of each protein was defined at the read of 0.5 on the fitting curve.

### Nanosecond and picosecond setups

Measurements were made using a time-resolved (TCSPC) miniTau fluorescence spectrometer (Edinburgh Instruments, Livingston, UK) in a 20 ns window divided into 2048 time channels. The fluorescence was excited using: (i) an EPL-450 picosecond laser (Edinburgh Instruments, Livingston, UK) with a central emission wavelength of 445.6 nm, pulse width (FWHM) of ∼90 ps@10 MHz driven at a repetition rate of 20 MHz; (ii) an EPLED-590 nanosecond pulsed LED (Edinburgh Instruments, Livingston, UK) with a central emission wavelength of 590 nm, pulse width (FWHM) of ∼1.3 ns driven at a repetition rate of 20 MHz. The photons were counted in the spectral range of 575-625 nm. The data processing, visualization and determination of χ2 (Pearson’s test) were carried out using the Fluoracle 2.5.1 software (Edinburgh Instruments, Livingston, UK).

### Fluorescence lifetime measurements

#### Femtosecond setup

The fluorescence decay kinetics of RFPs were recorded by a single-photon counting (SPC) detector with an ultra-low dark count rate (HPM-100-07C, Becker & Hickl, Germany), in the 620/10 spectral window, adjusted by an ML-44 monochromator (Solar, Belarus). Fluorescence was excited at 590 nm (repetition rate 80 MHz, pulse width 150 fs, optical power 5 mW) using the second harmonics (ASG-O, Avesta Project LTD.,Moscow, Russia) of a femtosecond optical parametric oscillator (TOPOL-1050-C, Avesta Project LTD.) pumped by a Yb femtosecond laser (TEMA-150, Avesta Project LTD.). The emission signal was collected perpendicular to the excitation beam. The temperature of the sample was stabilized during the experiment at 25 °C by a cuvette holder (Qpod 2e) with a magnetic stirrer (Quantum Northwest, USA). For the data acquisition, an SPCM Data Acquisition Software v. 9.89 (Becker & Hickl, Germany) was used. SPCImage software (Becker & Hickl, Germany) was used for the exponential fitting of fluorescence decays considering the incomplete decay of RFPs due to high repetition rate. The post-processing and visualization of the collected data were performed using an Origin Pro 2018 (OriginLab Corporation, USA).

### Photostability measurements

#### Purified proteins, low excitation intensity

For photobleaching experiments, the RFPs immobilized on TALON metal-affinity resin beads were imaged. Measurements were performed using a Leica laser scanning confocal inverted microscope DMIRE2 TCS SP2 (Leica Microsystems, Wetzlar, Germany) equipped with an HCX PL APO lbd.BL 63x 1.4NA oil objective and a 1.2 mW HeNe laser. Red fluorescent signal was acquired using the 543 nm excitation laser line and detected within 560-670 nm spectral range. The selected field of view (16x zoom) was scanned in a time-lapse (between frames) mode, wherein the sequence of detection and bleaching frames was repeated 500-1500 times without delay. To detect the red fluorescence signal, 10-20 % laser power and a PMT voltage of 700-800 V were used. To photobleach the fluorophores, 100 % laser power (giving about 2 W/cm^2^ power density) was used. The fluorescence data were all background-subtracted, averaged (n=5) and normalized to the maximum value. LaserCheck (Coherent, Saxonburg, Pennsylvania, USA) power meter was used to measure total power of the excitation light after the microscope objective. Light power density (W/cm^2^) was estimated by dividing the total power by the area of the laser-scanned region.

#### In cellulo measurement, high excitation intensity

For photobleaching experiments, the fluorescence signal of the RFPs, transiently expressed in the HeLa cell culture and lacked a specific intracellular targeting signal, was acquired. Measurements were performed using a Nanoimager S (ONI, Oxford, UK) microscope, equipped with an Olympus UPlanSApo x100 NA 1.40 oil immersion objective, 561 nm laser, 560 nm on-camera beam splitter and a Scope8 sCMOS camera. The cells were irradiated in epifluorescence mode with the 561 nm laser at a power density of 800 W/cm2 with simultaneous continuous signal recording and minimal delays between frames. Data analysis was performed using FiJi ImageJ 1.53f51 [28].

### Single-molecule localization microscopy

Super-resolution BALM imaging of the cytoskeleton of cultured mammalian cells was carried out as follows. Immediately before imaging, cell medium was changed to a minimal essential medium (MEM, Sigma-Aldrich, Saint Louis, Missouri, USA) supplemented with 20 mM HEPES. Single-molecule localization super-resolution imaging of living cells was performed using a Nanoimager S (ONI, Oxford, UK) microscope, equipped with an Olympus UPlanSApo x100 NA 1.40 oil immersion objective, 561 nm laser, 560 nm on-camera beam splitter and a Scope8 sCMOS camera. Imaging was performed using the following imaging condition set: 2 kW/cm^2^ 561 nm laser and 16.7 ms frame time (60 fps acquisition speed). Difference between signal-to-noise ratio of mKate2-K67R/R197H, TagRFP-T, and mKate2 localizations was Kolmogorov–Smirnov tested using test. Image acquisition and super-resolution reconstruction was performed using NimOS 490 1.18.3.15066 (ONI, Oxford, UK).

Image reconstruction was performed using default parameters. Data analysis was performed using FiJi ImageJ 1.53f51 [29] and custom Python 3.9 scripts.

## Results and discussion

To clarify the roles of particular amino acid substituents from the chromophore environment of FusionRed and mKate2 in determination of physicochemical distinctness of these fluorescent proteins (including their spectral differences and chromophore maturation peculiarities), we carried out a systematic mutational analysis implying an introduction of single, double and triple reciprocal substitutions (Fig. S1) at key positions 67, 158 and 197, which had earlier been identified as ‘gatekeepers’ of the FusionRed chromophore behavior based on its crystal structure [21].

### Description of the reciprocal mutants

#### Single mutations

Substitution at position 67 (Arg↔Lys) produced differential effects on the parental proteins. Thus, an mKate2-K67R variant was found to have negligible absorption in the visible range and to be almost non-fluorescent; the mutation probably strongly affected folding and/or chromophore maturation. Conversely, FusionRed-R67K possessed several well-marked spectral species, which likely correspond to different chromophore structures (Table 1, Fig. S2). Its absorption (being simultaneously a fluorescence excitation) peaked at 389, 514 and 580 nm. The latter red emissive species (*λ*_abs/ex_=580 nm, *λ*_em_=610 nm) behaves similar to the parental FusionRed, while both short wave forms are supposed to be the populations of immature chromophore. We assumed that a blue-emitting spectral form (*λ*_abs/ex_=389 nm, *λ*_em_=450 nm) corresponds to the neutral GFP-type chromophore, which is the well-described intermediate of the DsRed chromophore maturation [30, 31]. A yellow fluorescent form (*λ*_abs/ex_=514 nm, *λ*_em_=522 nm) of FusionRed-R67K, which spectrally resembles conventional yellow fluorescent proteins (EYFP, TagYFP) that bear a GFP-chromophore π-stacked with the tyrosine-203 residue [32], is less usual for RFPs. As one can speculate, R67K substitution led to a partial “freeze” of the FusionRed chromophore maturation at a pre-last oxidation step (GFP-like chromophore), and an anionic GFP-chromophore (usually absorbing at 470-500 nm) underwent a bathochromic spectral shift (to the yellow form) due to its π-stacking with the imidazole ring of histidine-197.

**Table 1.** Summary of the spectral properties, chromophore maturation and post-translational chemistry observed in the set of single, double and triple reciprocal mutants of FusionRed and mKate2.

| Protein | Absorption peak, nm | $\lambda_{ex}/\lambda_{em}$, nm | EC<sup>a</sup> (M<sup>-1</sup>cm<sup>-1</sup>) | FQY<sup>b</sup> | Molecular brightness (EC*QY/1000) | Comment |
| --- | --- | --- | --- | --- | --- | --- |
| FusionRed-R67K/C158A/H197R | 376; 488; 583 | 376/409; 488/508; 583/616 | n/d | <0.05 | n/d | bad maturation* |
| FusionRed-C158A/H197R | n/d | n/d | n/d | n/d | n/d | bad maturation* |
| **FusionRed-R67K/H197R** | 380; 488; 584 | 380/449; 488/511; 584/616 | n/d | 0.62 | n/d |  |
| FusionRed-R67K/C158A | 386; 513; 580 | n/d | n/d | n/d | n/d | bad maturation* |
| FusionRed- H197R | n/d | 570/607 | n/d | <0.01 | n/d | bad maturation* |
| **FusionRed-C158A** | 571 | 571/598 | 91 000 | 0.24 | 21.84 | [21] |
| **FusionRed-R67K** | 389; 514; 580 | 389/450; 514/522; 580/610 | n/d | 0.3 | n/d |  |
| **FusionRed** | **580** | **580/608** | **94 500** | **0.19** | **17.9** | [10] |
| **mKate2** | **586** | **588/633** | **62 500** | **0.4** | **25** | [20, 33] |
| mKate2-K67R | 405; 588 | n/d | n/d | n/d | n/d | bad maturation* |
| **mKate2-A158C** | 380; 590 | 590/624 | 47 300 | 0.47 | 22.2 | [21] |
| **mKate2-R197H** | 385; 510; 582 | 510/520; 582/612 | n/d | 0.26 | n/d |  |
| mKate2-K67R/A158C | n/d | n/d | n/d | n/d | n/d | bad maturation* |
| **mKate2-K67R/R197H** | **579** | **579/603** | **90 000** | **0.44** | **39.6** |  |
| **mKate2-A158C/R197H** | 380; 513; 583 | 380/435; 583/611 | n/d | 0.39 | n/d |  |
| mKate2-K67R/A158C/R197H | n/d | n/d | n/d | n/d | n/d | bad maturation* |

n/d – not determined
<sup>a</sup>Molar extinction coefficient; EC has not been determined for the variants possessing several spectral species;
<sup>b</sup>Fluorescence quantum yield; FQY was measured for the red emissive species only;
\*label applied if a low-to-invisible bacterial biomass fluorescence 48 h post transformation and/or low relative absorbance at the chromophore-related wavelengths (e.g., A280/A580 > 10) were observed.

An influence of the reciprocal mutation at position 158 (Cys↔Ala) on the spectral properties of FusionRed and mKate2 has earlier been documented [21]. Although this substitution did not result in formation of new spectral species and in severe inhibition of the chromophore maturation (or strong changes of molecular brightness), we include the data on the corresponding mutants (FusionRed-C158A and mKate2-A158C) to Table 1 for uniformity.

Mutations at position 197 (His↔Arg) showed effects in a way antagonistic to that of R67K/K67R. Similarly to mKate2-K67R, FusionRed-H197R was found to have a negligible absorption in the visible spectral region and to be almost non-fluorescent due to hindrance in the chromophore maturation. mKate2-R197H has the absorption maxima at 385, 510 and 582 nm, of which two latter are also the fluorescence excitation peaks, and emits at 520 and 612 nm (Table 1, Fig. S3). The identities and origins of these spectral forms are suggested to be the same as in FusionRed-R67K. Notably, there is a well-marked hypsochromic shift in absorption/emission maxima of the mKate2-R197H’s red form compared to the parental mKate2 protein (582/612 nm vs. 588/633 nm), that proves a key role of the His-197 in determination of the FusionRed spectral distinction.

#### Double mutations

The chromophore maturation in both proteins was generally less tolerant to an introduction of sets of two amino acid substitutions. Thus, the 3 of 6 double mutants were either extremely dim and weakly absorbing or almost non-fluorescent and having no detectable absorption maxima in the visible range (see table 1). In a relatively bright FusionRed-R67K/H197R variant, an antagonistic functionality of the residues at positions 67 and 197 is expressed. The R67K mutation partially unlocks the chromophore maturation strongly inhibited by the H197R. FusionRed-R67K/H197R possessed three emissive species (Table 1, Fig. S4): the blue-emitting neutral GFP (*λ*_abs/ex_=380 nm, *λ*_em_=449 nm), the green-emitting anionic GFP (*λ*_abs/ex_=488 nm, *λ*_em_=511 nm) and the red-emitting DsRed-like one (*λ*_abs/ex_=584 nm, *λ*_em_=616 nm). Remarkably, the red form of FusionRed-R67K/H197R showed a well-defined bathochromic shift in both absorption and emission (4 and 8 nm, respectively) compared to the parental FusionRed, thus giving an additional evidence on the role of the substituent at position 197 in the spectral tuning of the RFPs. The variant mKate2-A158C/R197H demonstrates complex spectral behavior (Table 1, Fig. S5), similar to that observed for the FusionRed R67K and mKate2 R197H proteins. In mKate2-A158C/R197H, the R197H substitution likely provides the hypsochromic shift of the red form’s fluorescence spectra as well as the stacking interaction with the immature “green” chromophore leading to formation of the spectral species with the absorption maximum at 513 nm. Importantly, the latter was found to be non-fluorescent.

mKate2-K67R/R197H was the only variant from the library of the mKate2/FusionRed reciprocal mutants which showed fast chromophore maturation and high molecular brightness. The fluorescence quantum yield of 0.44 and extinction coefficient of 90,000 make it 1.6 times brighter than the mKate2 and 2.2 times brighter than the FusionRed protein. In contrast to the parental protein, this double mutant lacks the minor shortwave absorption peaks at ∼390 and ∼450 nm attributed to immature chromophore and exhibits a blue-shifted main absorption band with a pronounced ‘shoulder’ at ∼540 nm, typical of FusionRed (Tables 1&2, Fig. 1).

**Table 2.**
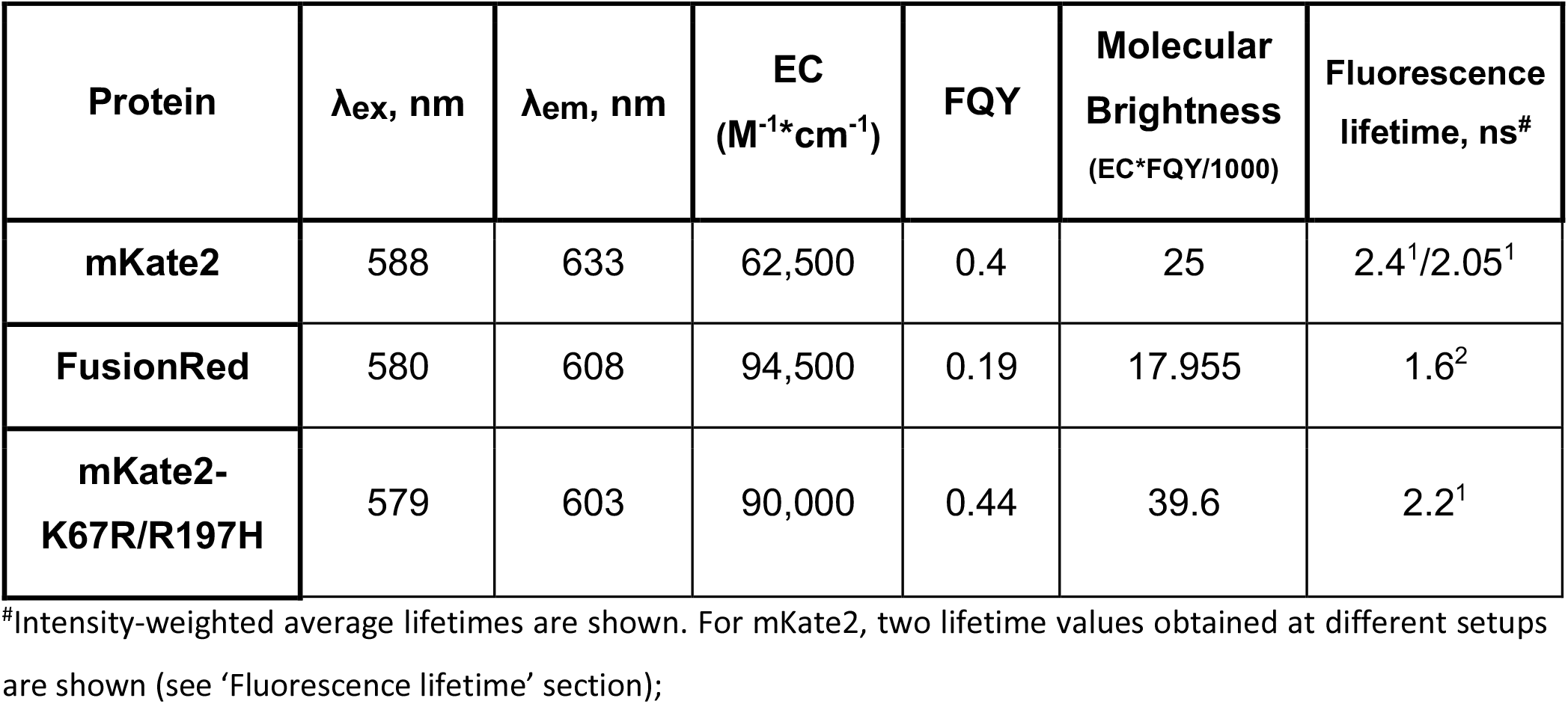
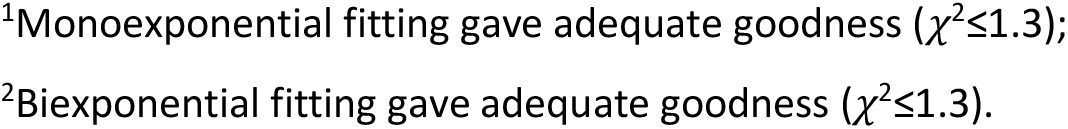
Brief summary of the spectral properties possessed by mKate2, FusionRed and mKate2-K67R/R197H aka Diogenes.

| Protein | $\lambda_{ex}$, nm | $\lambda_{em}$, nm | EC<br>($M^{-1} \cdot cm^{-1}$) | FQY | Molecular<br>Brightness<br>($EC \cdot FQY / 1000$) | Fluorescence<br>lifetime, ns<sup>#</sup> |
| --- | --- | --- | --- | --- | --- | --- |
| mKate2 | 588 | 633 | 62,500 | 0.4 | 25 | 2.4<sup>1</sup>/2.05<sup>1</sup> |
| FusionRed | 580 | 608 | 94,500 | 0.19 | 17.955 | 1.6<sup>2</sup> |
| mKate2-K67R/R197H | 579 | 603 | 90,000 | 0.44 | 39.6 | 2.2<sup>1</sup> |

<sup>#</sup>Intensity-weighted average lifetimes are shown. For mKate2, two lifetime values obtained at different setups are shown (see 'Fluorescence lifetime' section);
<sup>1</sup>Monoexponential fitting gave adequate goodness ( $\chi^2 \leq 1.3$ );
<sup>2</sup>Biexponential fitting gave adequate goodness ( $\chi^2 \leq 1.3$ ).

**Figure 1.**
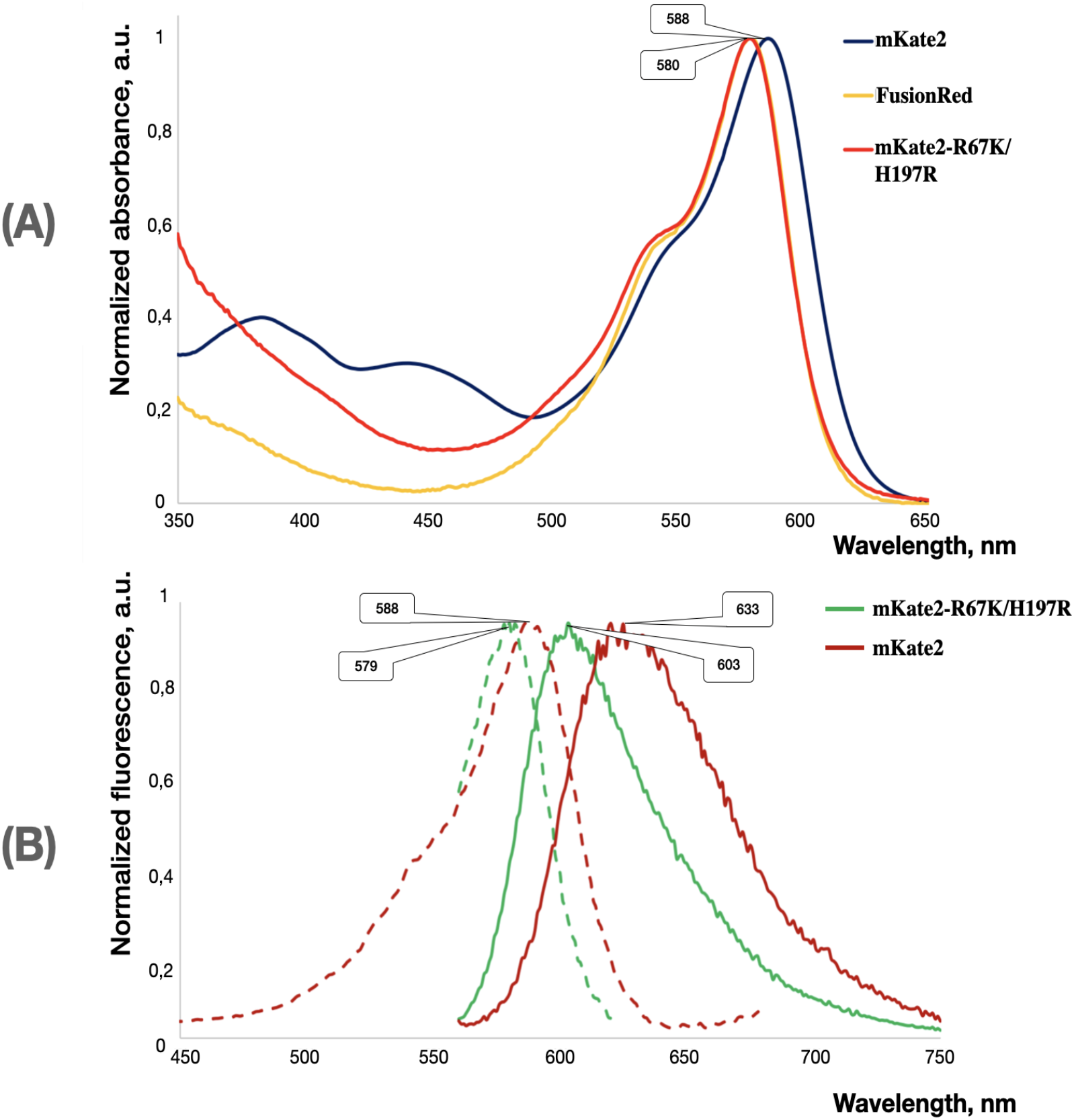
Absorption (**A**) and fluorescence (**B**) spectra of mKate2-K67R/R197H compared with those of its parent mKate2 and close homolog FusionRed (absorption only). Wavelengths of the major bands’ maxima are shown in the bubbles. In the fluorescence graph, dashed lines show fluorescence excitation, solid lines show fluorescence emission.

**Figure 2.**
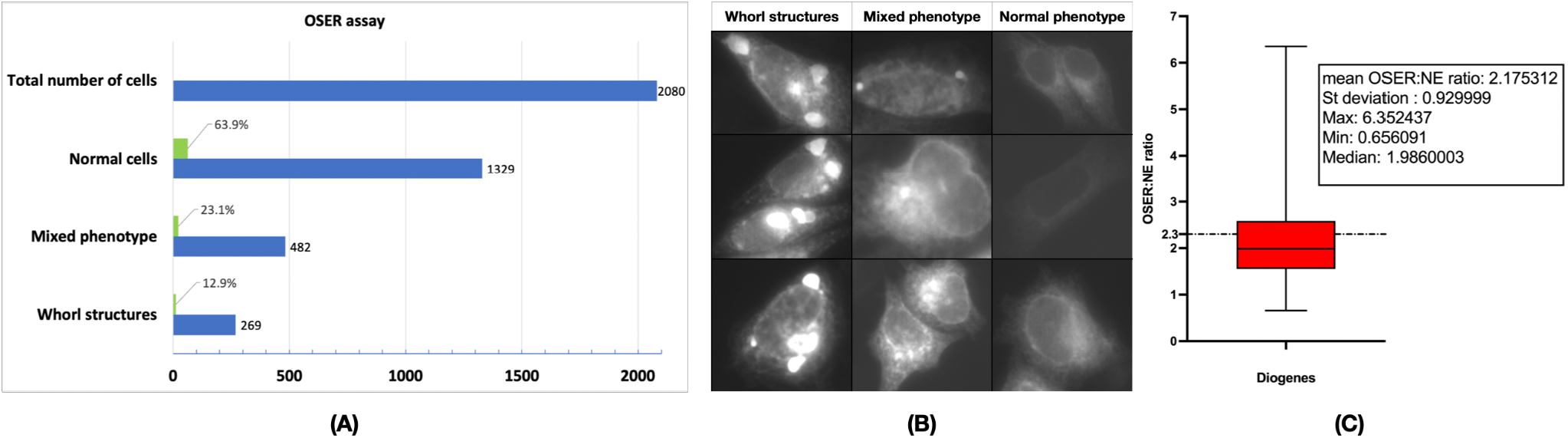
Summary of the Diogenes examination using OSER assay. (A) Histogram showing the total cell number (HeLa) and distribution of three phenotypes between them; (B) Gallery of the fluorescence images illustrating the phenotypes observed during the OSER-based examination in live HeLa cells. ‘Whorl structures’ row depicts the cells where a label’s homo-oligomerization led to a formation of the typical organized smooth endoplasmic reticulum (OSER) structures (whorls) as they have been earlier described [12, 23]. ‘Mixed phenotype’ row represents the cases when some labeling artifacts other than typical whorls (small puncta, dots, regions with locally increased brightness) are observed. ‘Normal phenotype’ row exemplifies the cells with evenly stained ER tubular network; (C) Graph showing the quantitative analysis of the OSER:NE intensity ratio in live U2OS cells. The red box indicates 25-75 percentile of the readings; whiskers stand for min and max read of the set. Median is illustrated by a horizontal line within the box; the dash-and-dot line at 2.3 indicates the monomer’s threshold OSER:NE value as according to the original paper [23]. Descriptive statistics for the data are shown in the inset.

#### Triple mutations

The introduction of the full triad of reciprocal 67/158/197 substitutions had a striking effect on the proteins’ maturation. Thus, both triple mutants – mKate2-K67R/A158C/R197H and FusionRed-C158A/H197R/R67K – displayed undetectable to very low absorbance/fluorescence in the visible part of the spectrum (see Table 1), presumably indicating the “freezing” of the chromophore in the early stages of maturation. The informativeness of these variants in terms of establishing the molecular determinants of the spectral distinctiveness of FusionRed/mKate2 turned out to be low.

Generally, our phenotypic analysis revealed that the substitutions at position 67 likely cause a shift of the chromophore within the beta-barrel since after the mutagenesis, the spectroscopic signs of its maturation process.

pi-stacking interaction with histidine-197 (if any) become changed compared to those in the parental proteins. The influence of these substitutions on the chromophore maturation is also evident: in all cases except the mKate2-K67R/R197H they led to either a strong maturation alteration or at least an elevated presence of the shortwave spectral species representing the maturation intermediates. Substitutions at position 197 induce spectral shifts, a bathochromic absorption/emission shift in the case of the FusionRed-derived variants and a hypsochromic one in the mKate2 mutants. We suppose that the reason for this phenomenon might be connected with the π-stacking interaction between the chromophore and histidine (occupying position 197 in the original FusionRed) switched off/on by reciprocal mutations. Additionally, these mutations demonstrate a noticeable impact on the chromophore

### Physicochemical properties of mKate2-K67R/R197H and its performance in microscopy

We next aimed at a detailed characterization of physicochemical properties of the mKate2-K67R/R197H protein, named Diogenes, focusing on its performance in cellular fluorescence imaging.

#### Oligomeric state and protein labeling

The first step was to analyze the protein’s oligomeric state that is among the key predictors of efficient low-disturbed/minimally invasive labeling of intracellular targets. According to the gel-filtration chromatography data (Fig. S6, Fig. S7), the purified protein elutes as a single peak with an estimated molecular weight of ∼38 kDa (at a concentration up to at least 5 mg/ml). Since this molecular weight corresponds neither a monomer (∼25 kDa) nor a dimer (∼50 kDa), the gel-filtration result cannot be interpreted unambiguously. One can assume that concentrated Diogenes in aqueous solution is either a strict monomer or a strict dimer, having anomalous chromatographic mobility in both cases. Alternatively, it is possible that we observed an equilibrium mixture of the monomeric and dimeric states.

It would be reasonable to expect that in terms of oligomeric state, Diogenes will be close to the parental mKate2, which was originally described as a monomer [20], with further evidence of some propensity to oligomerize in aqueous solutions at high concentration [10] and in cellulo [12]. At the same time, it is important to evaluate how the monomeric quality of this variant compares with that of its spectral analog FusionRed. During the engineering of the latter, considerable effort was devoted to optimizing the outer surface of the beta-barrel, including the elimination of potentially dimerizing residues [10]. It was indeed shown that purified FusionRed behaves as a strict monomer [10], and gets a higher monomericity rank than mKate2 when examined in cells (91.5 ± 3.0% vs. 81.1 ± 6.1% in the OSER assay [12]).

However, the causal link between the monomerizing mutations introduced to FusionRed and its better performance in cellulo remains somewhat debatable, as the rational design of these substitutions was based not on the spatial structure of mKate2, but on that of mKate [34]. Moreover, protein folding and observed molecular interactions in crystals may not fully correspond to those in aqueous phase [35, 36]. In any case, the ambiguous chromatographic picture for Diogenes prompted us to evaluate its oligomeric state in a cellular model system.

To this end, we applied an OSER assay [23], which has become a de facto standard for assessing the monomerization of fluorescent proteins in cellulo [12, 37]. Due to modern uses of OSER assay oftenly diverging from the original one, we performed it in two different cell lines - HeLa for the widely-used simplified (OSER to non-OSER phenotypic) assessment and U2OS for the original (OSER:NE ratio) assessment. In HeLa, the analysis revealed ∼87% whorl-free cells (Fig. S7), that could be interpreted as a relatively high monomeric quality and is in between the FusionRed and mKate2 scores published previously [12]. It should be mentioned, however, that additionally to obvious OSER-negative/positive cells we observed a well-represented (∼23%) cell fraction possessing diverse labeling features, such as small puncta, dots, local areas with increased brightness), which probably should not be attributed as a typical tubular ER phenotype (we labeled this population ‘mixed phenotype’, see Fig. S7 for details). Structures mentioned above can indicate protein aggregation or its non-specific interactions with the intracellular environment, which could probably limit its efficiency in some circumstances. As according to the original OSER analysis protocol in U2OS, the revealed mean OSER:NE ratio of Diogenes is 2.175 with median of 1.986 and standard deviation of 0.9299. Despite the relatively large standard deviation, both the mean OSER:NE ratio and the median allow us to consider Diogenes monomeric - with the monomericity borderline set as OSER:NE ≤2.3 ± 0.6 [23].

Finally, we assembled several mammalian expression constructs for visual evaluation of the effectiveness of mKate2-K67R/R197H when working in fusions. For this testing, we selected targets (cytoskeleton proteins) whose visualization quality, according to our experience, significantly depends on the oligomeric status of the tag (Fig. 3). Subjectively, we rate the labeling quality as very high.

**Figure 3.**
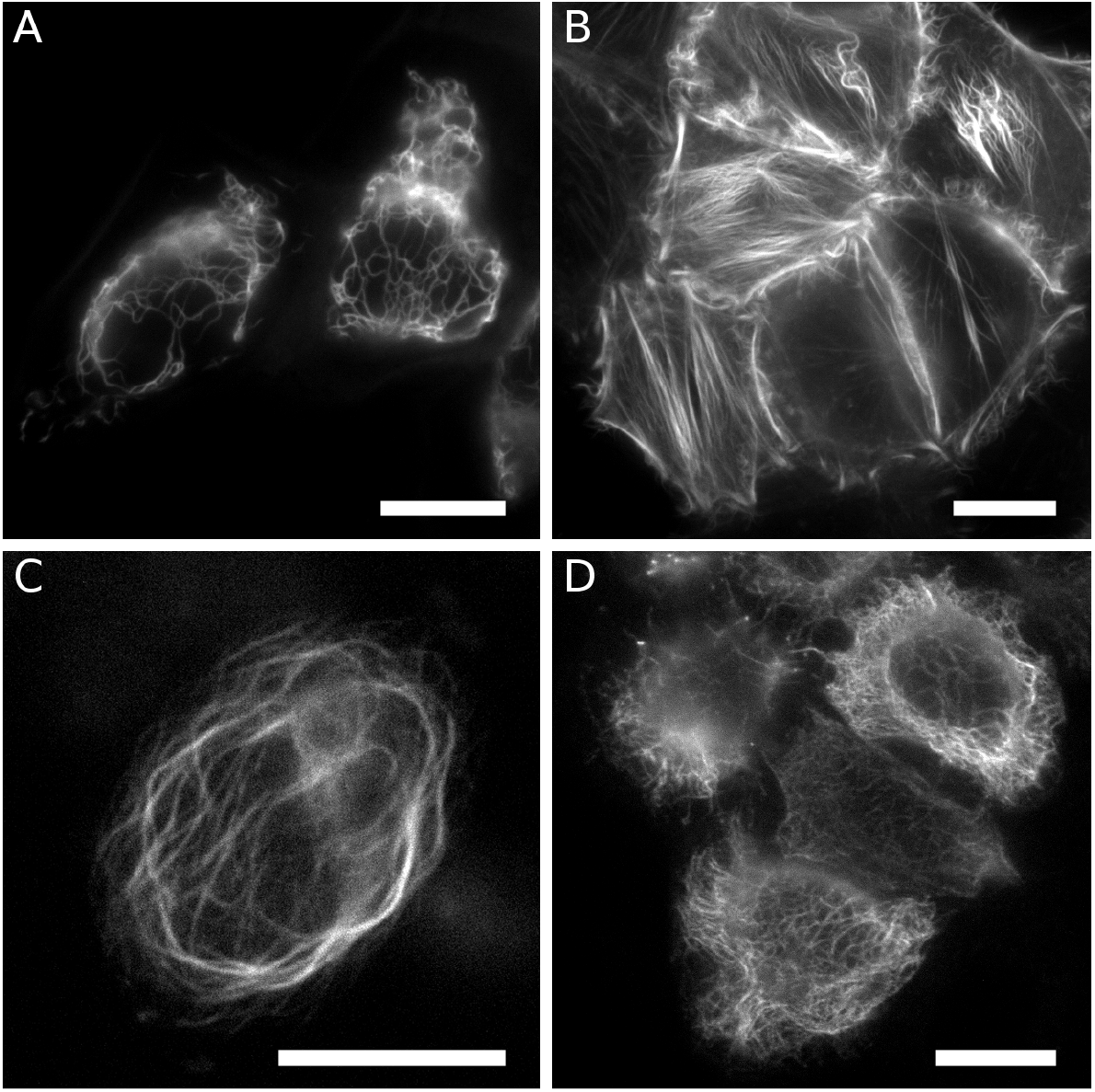
Fluorescent labeling of intracellular structures with Diogenes in live HeLa Kyoto cells. Vimentin-Diogenes (**A**), lifeact-Diogenes (**B**), ensconsin-Diogenes (**C**), Diogenes-cytokeratin (**D**); Scale bars are 15 μm.

#### pH-stability

We next compared a stability of fluorescence intensity between Diogenes and its relatives, mKate2 and FusionRed, within the wide pH range of 3-11 (Fig. 4, S8). Generally the protein exhibited high pH-stability, similar to that of mKate2, which is among the most pH-stable RFPs. Specifically, it could maintain a fluorescence level of ≥80% of the maximum within the most physiologically and biochemically relevant pH range of 6.5-9.5. In the acidic range (pH 3-6), Diogenes showed worse relative brightness than FusionRed but slightly surpassed mKate2 (their pKa values were determined to be 6.1, 5.76 and 6.16, respectively). Unlike both counterparts, the Diogenes fluorescence sharply decreases in the strongly alkaline pH range of 10-11 though this acidity level is not much biologically relevant.

**Figure 4.**
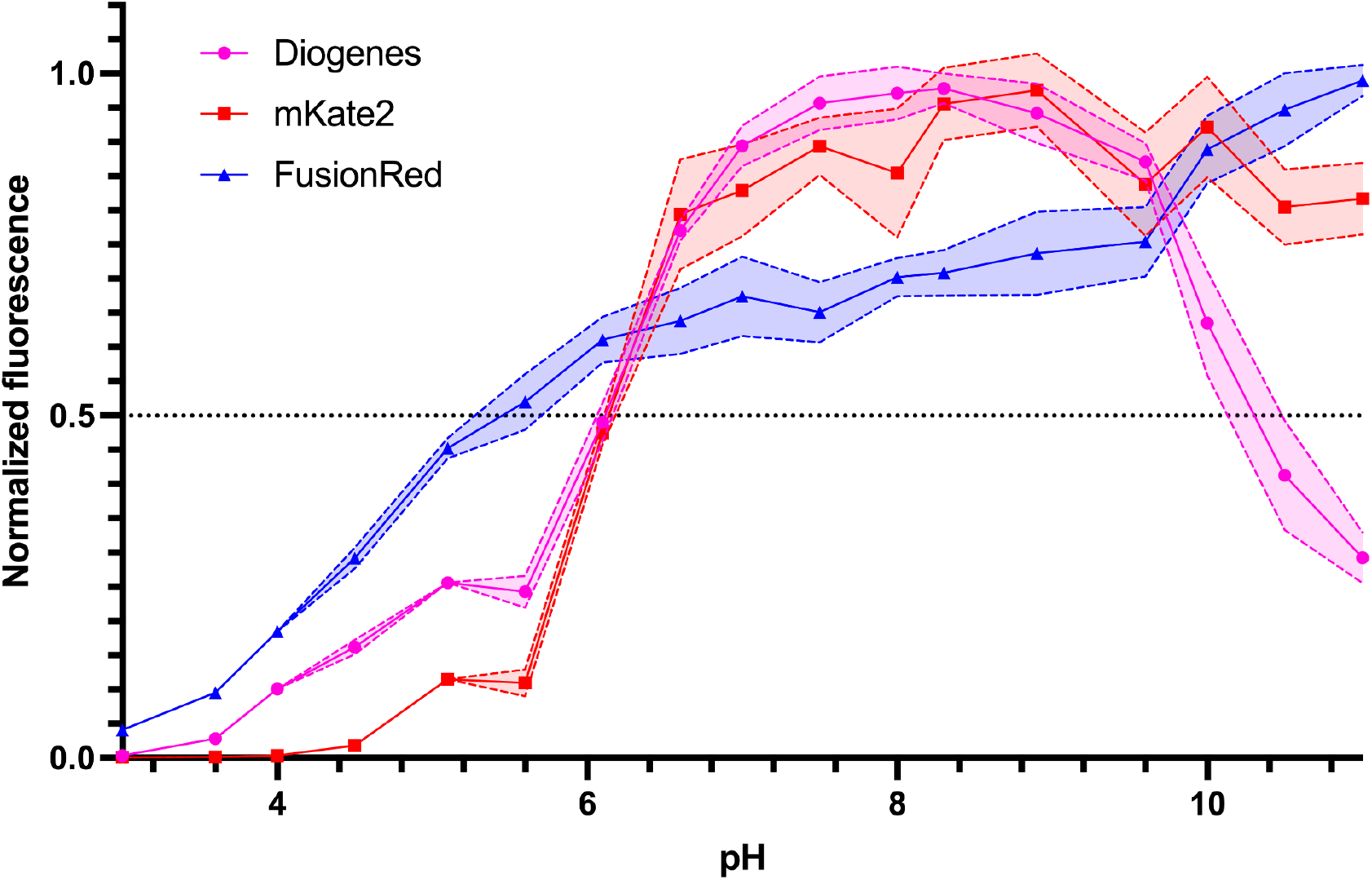
Graph showing the fluorescence intensity dependence on pH (pH-stability) measured for the purified mKate2-K67R/R197H (Diogenes), mKate2 and FusionRed proteins. Signal at each measurement point was normalized to a maximum signal value within the dataset.

#### Fluorescence lifetime

We then measured fluorescence decay kinetics of the purified mKate2-K67R/R197H (Diogenes) in aqueous solution using the time-correlated single photon counting approach and 3 different instrument set-ups (Fig. 5, Figs. S9 & S10). Importantly, the decay was shown to be monophasic in all cases, with a lifetime value of ∼2.2 ns. Surprisingly, in contrast to it and the FusionRed protein, which showed a biphasic fluorescence decay and a mean lifetime of ∼1.6 ns with every set-up used, parental mKate2 displayed a noticeable dependence of its lifetime on the measurement equipment. Thus, upon excitation with a 450 nm picosecond laser (FWHM ∼100 ps, 20 MHz) and a 590 nm nanosecond pulsed LED (FWHM ∼1.5 ns, 20 MHz) its lifetime was 2.4 ns (see Fig. S9 and Fig. 5), while with a 590 nm femtosecond laser (FWHM ∼150 fs, 80 MHz) – only 2.05 ns (Fig. S8). The reasons for such flexibility remain unclear; it might be connected with some kind of excited-state processes known to occur in mKate2 and related proteins [30, 38].

**Figure 5.**
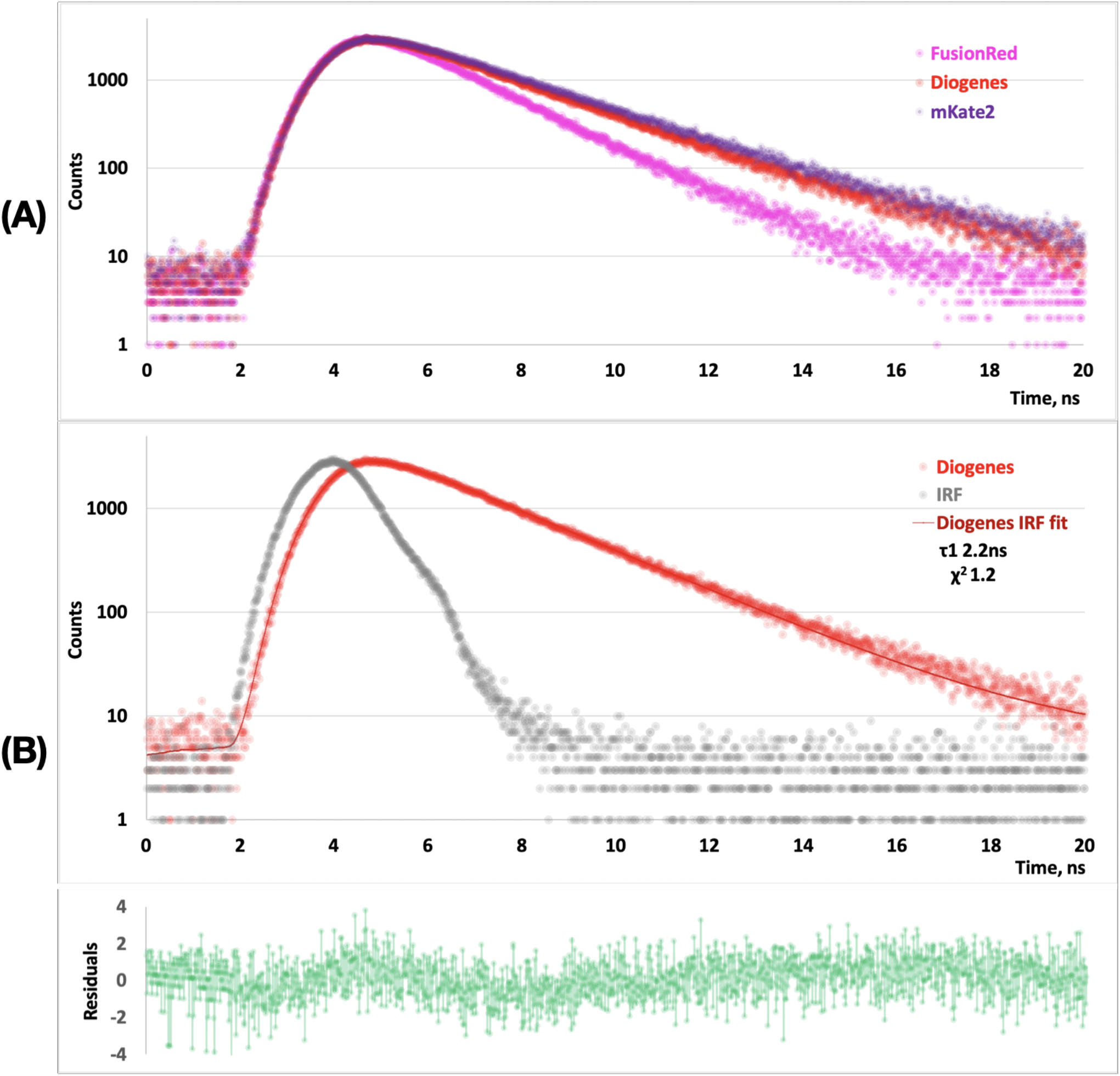
Fluorescence decay kinetics recorded upon 590 nm single-photon excitation with the nanosecond pulsed LED driven at a repetition rate of 20 MHz. Comparison of the raw-data decay curves for FusionRed, mKate2 and Diogenes (**A**). Single-component exponential fitting of the Diogenes decay curve (**B**). Deconvolution with IRF was used for fitting. Measured instrument response (IRF) is shown in red.

Taking into account the excitation/emission wavelengths of Diogenes, mRuby [39] or mRuby2 [40] could be considered as its close competitors in terms of fluorescence brightness/lifetime.

#### Photostability

High photostability is a desirable fluorophore property for both conventional fluorescence imaging and advanced microscopy techniques [41]. Moreover, photobleaching rate of fluorescence protein may depend non-linearly on the excitation source power [12, 41]; this phenomenon can have a considerable impact when choosing a specific probe variant for a particular experiment. In this regard, we measured the photostability of Diogenes in two different model systems (see Fig. 6). The photostability of purified mKate2-K67R/R197H measured in an aqueous environment (protein immobilized on microparticles) at a moderate power density typical for widefield fluorescence microscope (∼2 W/cm^2^) was found to be slightly higher than that of FusionRed (bleaching t_1/2_ 215 s vs. 165 s) and significantly lower than that of mKate2 (t_1/2_ ∼590 s, Fig. 6A). Surprisingly, in live HeLa cells upon high-intensity (∼kW/cm^2^) excitation, typical for SMLM techniques, the new RFP showed better performance than mKate2 (circa twofold higher photostability, Fig. 6B) and approximately the same one as FusionRed, which is much dimmer and was therefore expected to be more photostable.

**Figure 6.**
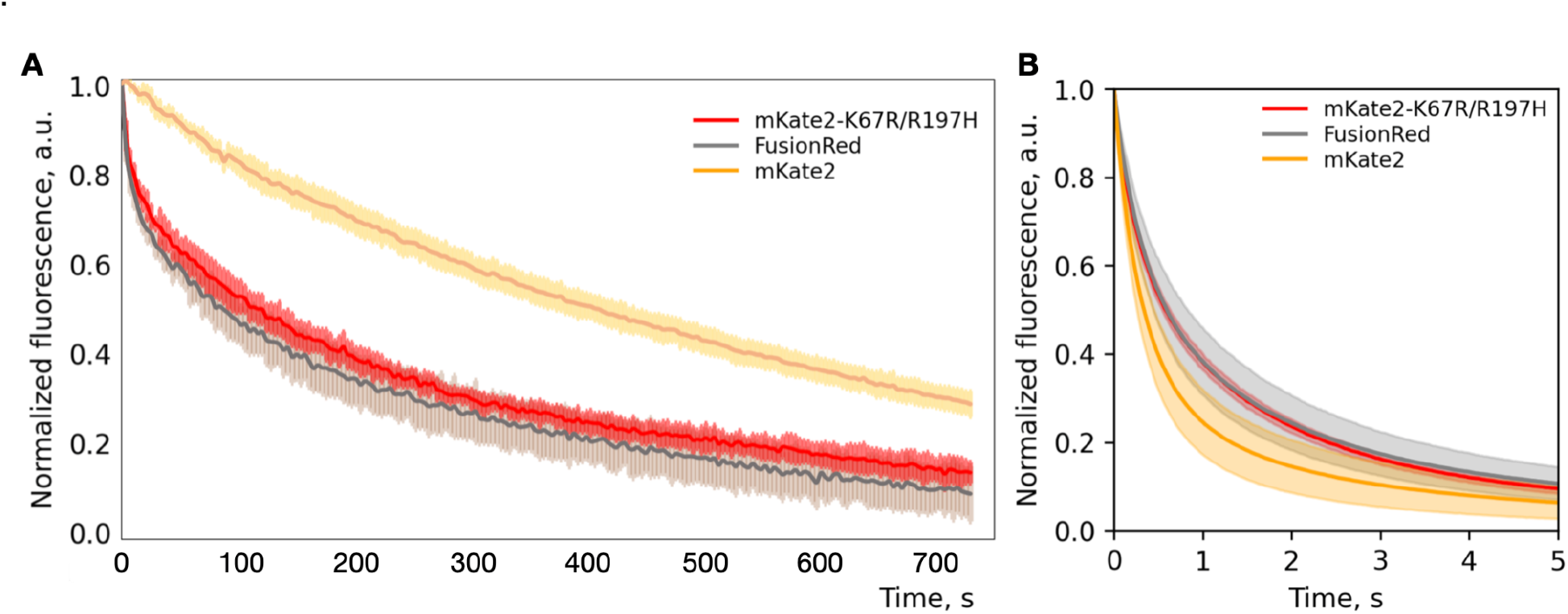
Photobleaching kinetics of the red fluorescent proteins mKate2-K67R/R197H (aka Diogenes), mKate2 and FusionRed measured in aqueous solution of the purified protein at an excitation power density ∼2 W/cm^2^ (**A**) and live HeLa cells at ∼1 kW/cm^2^ (**B**). Solid lines indicate mean fluorescence intensity during photobleaching. Transparent areas indicate standard deviation (5 protein-containing particles or 20 cells for each fluorescent protein).

### Single-molecule behavior of mKate2-K67R/R197H

The increased photostability of Diogenes observed during imaging in a high excitation power mode, similar to that used in single-molecule microscopy techniques, prompted us to investigate the protein’s behavior at the single-molecule level. Preliminary runs performed using dSTORM-like settings of the superresolution fluorescence microscope on droplets of the purified protein revealed pronounced stochastic blinking behavior of Diogenes (data not shown). Critically, red and far-red FPs, including variants such as mScarlet, mKate2, TagRFP, FusionRed, and FusionRed-MQ, although exhibiting blinking behavior [28, 42, 43], mostly fall short of green fluorescent proteins in terms of single-molecule performance, with only a few exceptions [42]. Therefore, evaluating the potential of new RFP variants for various SMLM techniques, where spontaneous fluorescence blinking can be utilized to refine localization of labeled molecules, is of importance.

Here we applied Single-Molecule Localization Microscopy (SMLM) to reveal whether the Diogenes variant is capable of spontaneous blinking in cellulo and visualizing intracellular structures with enhanced resolution. We took the parental protein mKate2 and TagRFP-T, a protein known to have the strongest blinking pattern among the previously examined RFPs [42], as references. Comparison of the single-molecule performance of Diogenes, TagRFP-T, and mKate2 was carried out in a model system where these probes were fused with vimentin, transiently expressed in live HeLa cells and monitored under super-resolution microscopy conditions (Fig. 7).

**Figure 7.**
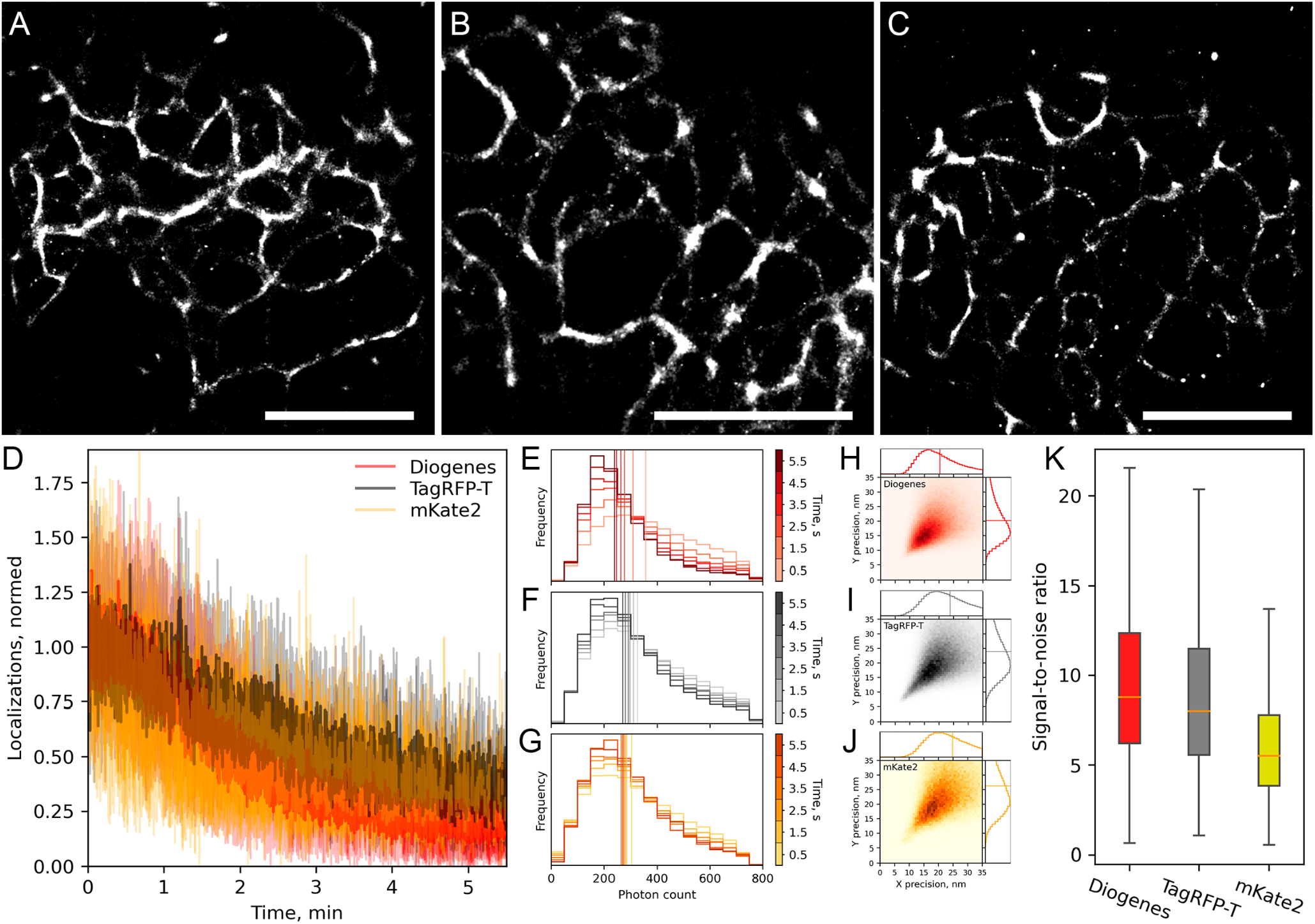
Comparison of live-cell super-resolution imaging performance of mKate2-K67R/R197H, TagRFP-T, and mKate2 as parts of vimentin fusion proteins in live HeLa cells under the following imaging conditions: 2 kW/cm^2^ 561 nm laser, 16.7 ms frame time, 20,000 frames. Super-resolution images of live HeLa cells transfected with vimentin-Diogenes, vimentin-TagRFP-T, and vimentin-mKate2 respectively (**A**,**B**,**C**); Scale bars are 5 μm. Stability of localization density of Diogenes, TagRFP-T, and mKate2 (**D**). Histogram of changes in the number of detected photons per single-molecule event over time of Diogenes, TagRFP-T, and mKate2 respectively; vertical lines represent median values (**E**,**F**,**G**). 2D histograms of localization precision per single-molecule event of Diogenes, TagRFP-T, and mKate2 respectively (**H**,**I**,**J**); vertical lines on 1D histograms represent median values. Signal-to-noise ratio of detected localizations (**K**); whiskers show standard deviation, orange horizontal lines indicate median values.

Under the illumination of 2 kW/cm^2^ at 561 nm light and with a 16.7 ms frame time, all proteins blinked at a single molecule level, allowing the reconstruction of the subdiffraction image of vimentin fibers in live HeLa cells (Fig. 7A-C). Comparative analysis revealed that the stability of localization density was similar for all three proteins (Fig. 7D). Additionally, the difference in the median single-molecule brightness value was negligible among these proteins (Fig. 7E-G). The localization accuracy of Diogenes appeared to be slightly better than that of mKate2 and TagRFP-T (20.5 nm vs 24 nm and 25 nm, respectively, Fig. 7H-J), while the median value of the signal-to-noise ratio was ∼ 1.5-fold higher for Diogenes than for mKate2, though the differences in the overall datasets for signal-to-noise values were insignificant (Fig. 7K). In conclusion, Diogenes’ performance in live-cell SMLM is on par or even slightly better than that of RFP, which was previously described as a promising probe for this microscopy modality [42]. However, it is still inferior in terms of stability of localization density, molecular brightness, and localization precision to green fluorescent proteins capable of spontaneous blinking (e.g. mNeonGreen [44] and mBaoJin [45]).

Intriguingly, additional UV illumination during imaging affected the density of localizations of all three proteins (Fig. S10). Short UV laser spikes significantly increased the number of recorded localizations of all three proteins. Although this experiment is not sufficient to conclude about the nature of this phenomenon, it can be assumed that UV illumination may induce additional maturation of the chromophore or switch the chromophore from a long-lived dark state to a fluorescent state.

## Conclusions

In this study, we systematically inspected the library of reciprocal mutants of the far-red fluorescent protein mKate2 and its daughter, the red FusionRed. We aimed to clarify the particular role of three residues from the chromophore environment (Arg/Lys-67, Cys/Ala-158, His/Arg-197) in determining the photophysical identities of these widely used genetically encoded probes.

One of the representatives of the constructed library, mKate2-K67R/R197H, which we named Diogenes, demonstrated a good combination of physicochemical and spectral properties and represents a promising probe for conventional and advanced fluorescence microscopy techniques. It inherits the advantages of both related proteins (FusionRed and mKate2). In particular, it has high fluorescence brightness, demonstrates a monophasic fluorescence decay like mKate2, and shows good performance as a fusion tag like FusionRed. In terms of monomerization, Diogenes surpasses the parental mKate2 and possibly approaches the monomeric quality of FusionRed. It is worth noting the relatively high photostability of Diogenes (especially when normalized to the molecular brightness of the protein) under conditions of intense irradiation, as well as its remarkable ability for UV-induced photoactivation, which likely opens up possibilities for modulating its single-molecule behavior under live-cell multiphoton microscopy. Our results include indirect evidence that a smaller fraction of molecules trapped in long-lived transient dark states might be present in the population of Diogenes molecules (Fig. S10). Together with the data of absorption spectroscopy (Fig. 1A), this may indicate a higher quality of chromophore maturation and its steric adaptation inside the protein molecule compared to related RFPs.

It is important to note that compared to its spectral analog, FusionRed, Diogenes carries a minimal number of mutations relative to the parental mKate2.

Furthermore, the combination of substitutions (K67R/R197H) we found in the reciprocal library analysis was previously independently transferred from the bright but oligomerization-prone TagRFP protein to the dim monomer mKate2.5 to obtain FusionRed [10]. In this regard, it would be interesting to compare high-resolution structural data along with molecular dynamics (preferably high-resolution NMR data) for the FusionRed and Diogenes pair, to decipher the roles played by the peripheral amino acid residues that differentiate these proteins. Like its relative FusionRed, Diogenes can become a template for future semi-rational optimizations of RFPs, including those using high-throughput approaches.

## Supporting information

All supplementary figures and tables for the manuscript

## Supporting information

Additional figures demonstrate the scheme of reciprocal FusionRed-mKate2 mutagenesis procedure (Figure S1); absorbance and fluorescence spectra of FusionRed-K67R (Figure S2), mKate2-R197H (Figure S3), FusionRed-R67K/H197R (Figure S4), mKate2-A158C/R197H (Figure S5); gel filtration/liquid chromatography (Figure S6) and its calibration plot with linear fit (Figure S7) with molecular weights of proteins standards used in calibration (Table S1); mKate2, FusionRed and Diogenes pH stability sigmoidal fits (Figure S8), mKate2, FusionRed and Diogenes fluorescence decay kinetics upon 590 nm and 450 nm single-photon excitation (Figures S9 and S10, respectively); UV-laser illumination effect on localization density of mKate2-K67R/R197H, TagRFP-T, and mKate2 (Figure S11).

## Author contributions

**Conceptualization:** Dmitry A. Ruchkin, Vladimir Z. Pletnev, Alexey M. Bogdanov

**Data Curation:** Dmitry A. Ruchkin, Alexey S. Gavrikov, Andrey Yu. Gorokhovatsky, Eugene G. Maksimov, Vladimir Z. Pletnev, Alexey M. Bogdanov

**Formal Analysis:** Dmitry A. Ruchkin, Alexey S. Gavrikov

**Funding Acquisition:** Nadya V. Pletneva

**Investigation:** Dmitry A. Ruchkin, Alexey S. Gavrikov, Danila V. Kolesov, Andrey Yu. Gorokhovatsky, Tatiana V. Chepurnykh, Eugene G. Maksimov, Nadya V. Pletneva, Anastasiia M. Pavlova, Vladimir A. Nikitin, Alexey M. Bogdanov

**Methodology:** Dmitry A. Ruchkin, Alexey S. Gavrikov, Danila V. Kolesov, Andrey Yu. Gorokhovatsky, Alexander S. Mishin, Vladimir Z. Pletnev, Alexey M. Bogdanov

**Project Administration:** Alexey M. Bogdanov

**Resources:** Andrey Yu. Gorokhovatsky, Alexander S. Mishin, Vladimir Z. Pletnev, Konstantin A. Lukyanov, Alexey M. Bogdanov

**Supervision:** Danila V. Kolesov, Konstantin A. Lukyanov, Alexey M. Bogdanov

**Visualization:** Dmitry A. Ruchkin, Alexey S. Gavrikov, Andrey Yu. Gorokhovatsky

**Writing – Original Draft Preparation:** Alexey M. Bogdanov, Dmitry A. Ruchkin, Alexey S. Gavrikov

**Writing – Review & Editing:** Alexey M. Bogdanov, Dmitry A. Ruchkin The manuscript was written through contributions of all authors. All authors have given approval to the final version of the manuscript.

## Protein accession IDs

FusionRed (NCBI Accession: 6U1A_A); mKate2 (NCBI Accession: AEX25288.1); Cytochrome p450 for CytERM-based (OSER) examination (NCBI Accession: KAF4413803.1); Vimentin [Homo sapiens] (NCBI Accession: NP_003371.2); Lifeact (NCBI Accession: 7AD9_G); Ensconsin [Homo sapiens] (NCBI Accession: NP_001375262.1); Cytokeratin 18 [Homo sapiens] (NCBI Accession: CAA31375.1); TagRFP-T (NCBI Accession: 5JVA_A).

## Acknowledgement

This study was supported by a Grant from the Russian Science Foundation (RSF), project № 23-24-00011.

