## Supplementary material for "The two key substitutions in the chromophore environment of mKate2 to produce an enhanced FusionRed-like red fluorescent protein": All supplementary figures and tables for the manuscript

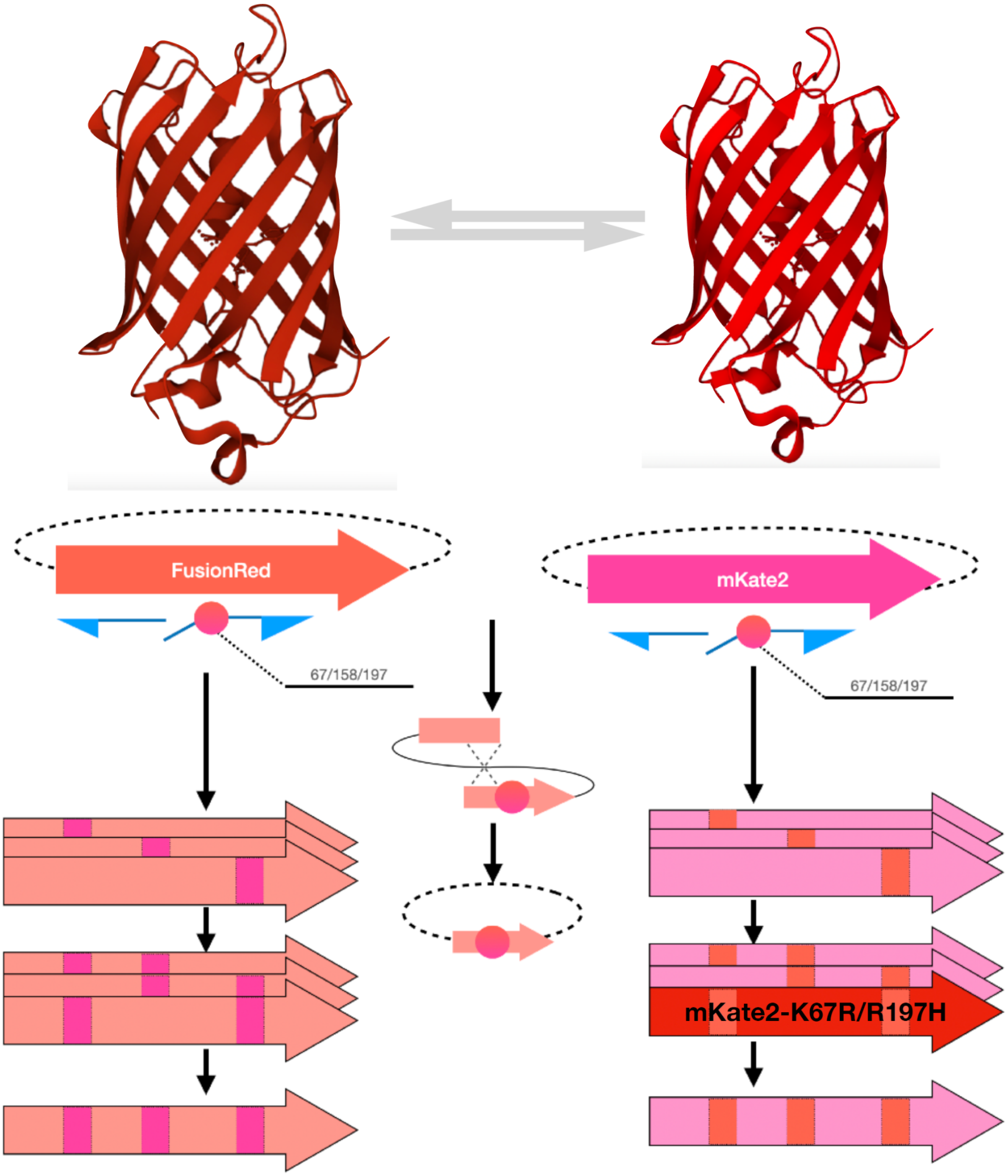


**Supplementary Figure S1**. Scheme showing how the library of point reciprocal mutants used in this study was engineered. The upper panel illustrates a general idea of 'exchanging' amino acid residues at positions 67/158/197 between the original FusionRed and mKate2 proteins. In the center of the scheme, there is a principle of IVA-cloning-based site-directed mutagenesis explained graphically (blue arrows are oligonucleotides, pink circles are DNA-mismatches with substituted codons, dashed-lined ellipses depict vector backbone, dotted cross shows recombination and ligation of full-vector-PCR product). Aligned thick arrows on the left and right represent particular variants carrying single, double and triple (single arrow) substitutions (mutated sites are shown as vertical bands of the 'opposite' color).

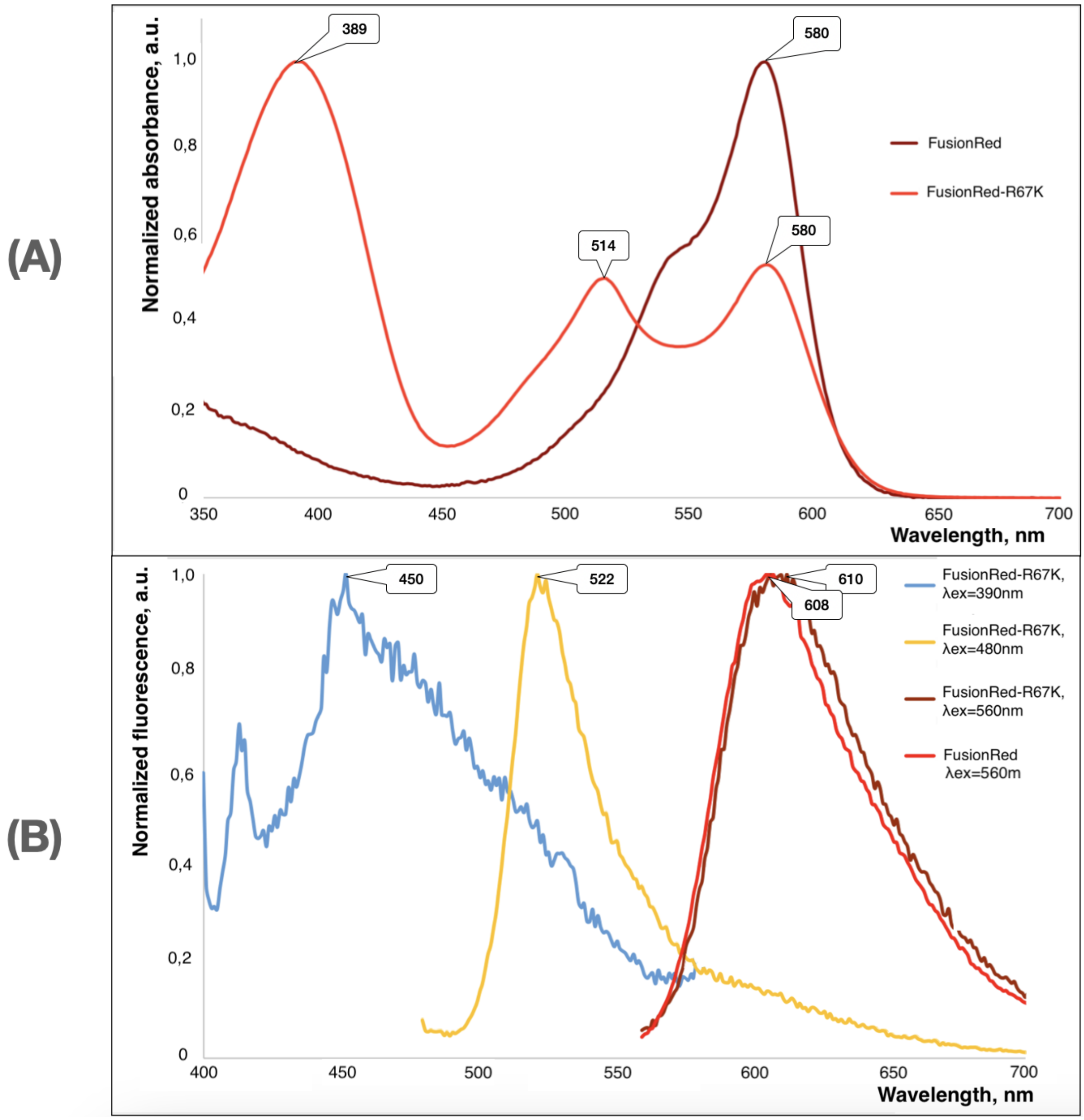


**Supplementary Figure S2.** Absorption (A) and fluorescence emission (B) spectra of FusionRed-R67K compared with those of its parent FusionRed. Wavelengths of the major bands' maxima are shown in the bubbles. In the fluorescence graph, excitation wavelengths used for the emission spectra recording are shown in the legend (right upper corner).

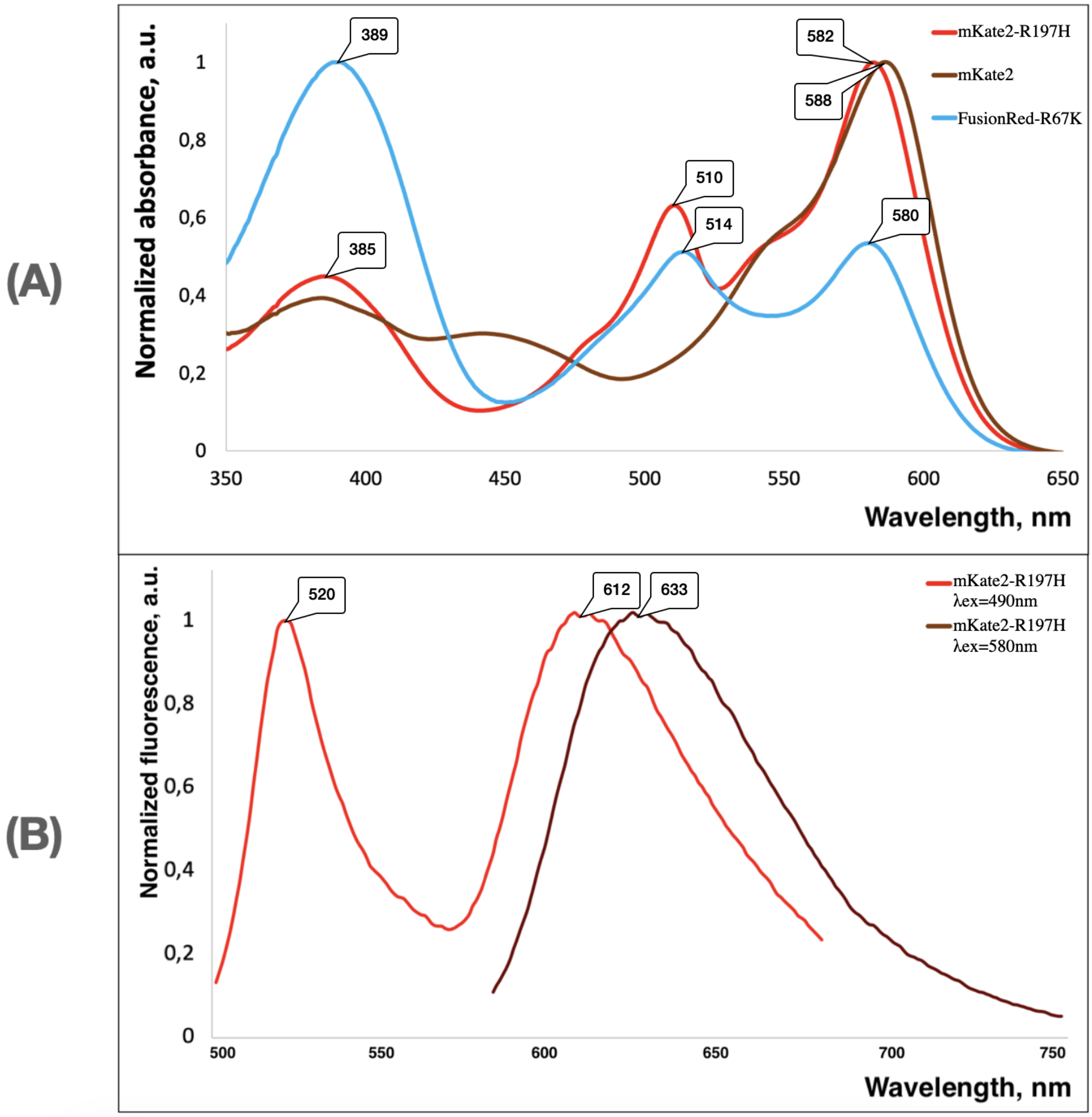


**Supplementary Figure S3.** Absorption (A) and fluorescence emission (B) spectra of mKate2-R197H compared with those of its parent mKate2. Wavelengths of the major bands' maxima are shown in the bubbles. In the absorption graph, the spectrum of FusionRed-R67K (solid blue line) is also added as a reference. In the fluorescence graph, excitation wavelengths used for the emission spectra recording are shown in the legend (right upper corner).

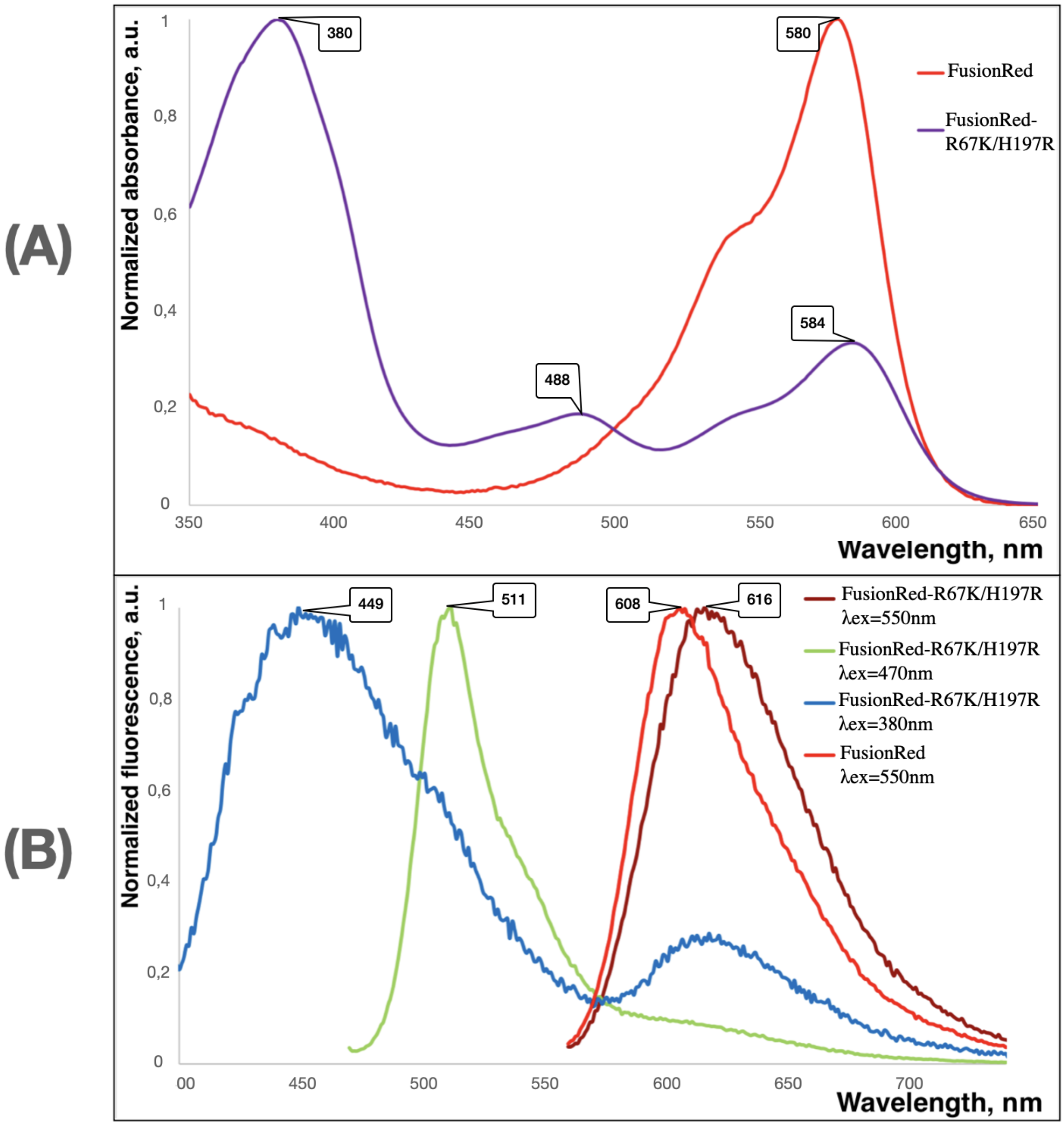


**Supplementary Figure S4.** Absorption (A) and fluorescence emission (B) spectra of FusionRed-R67K/H197R compared with those of its parent FusionRed. Wavelengths of the major bands' maxima are shown in the bubbles. In the fluorescence graph, excitation wavelengths used for the emission spectra recording are shown in the legend (right upper corner).

(A)

(B)

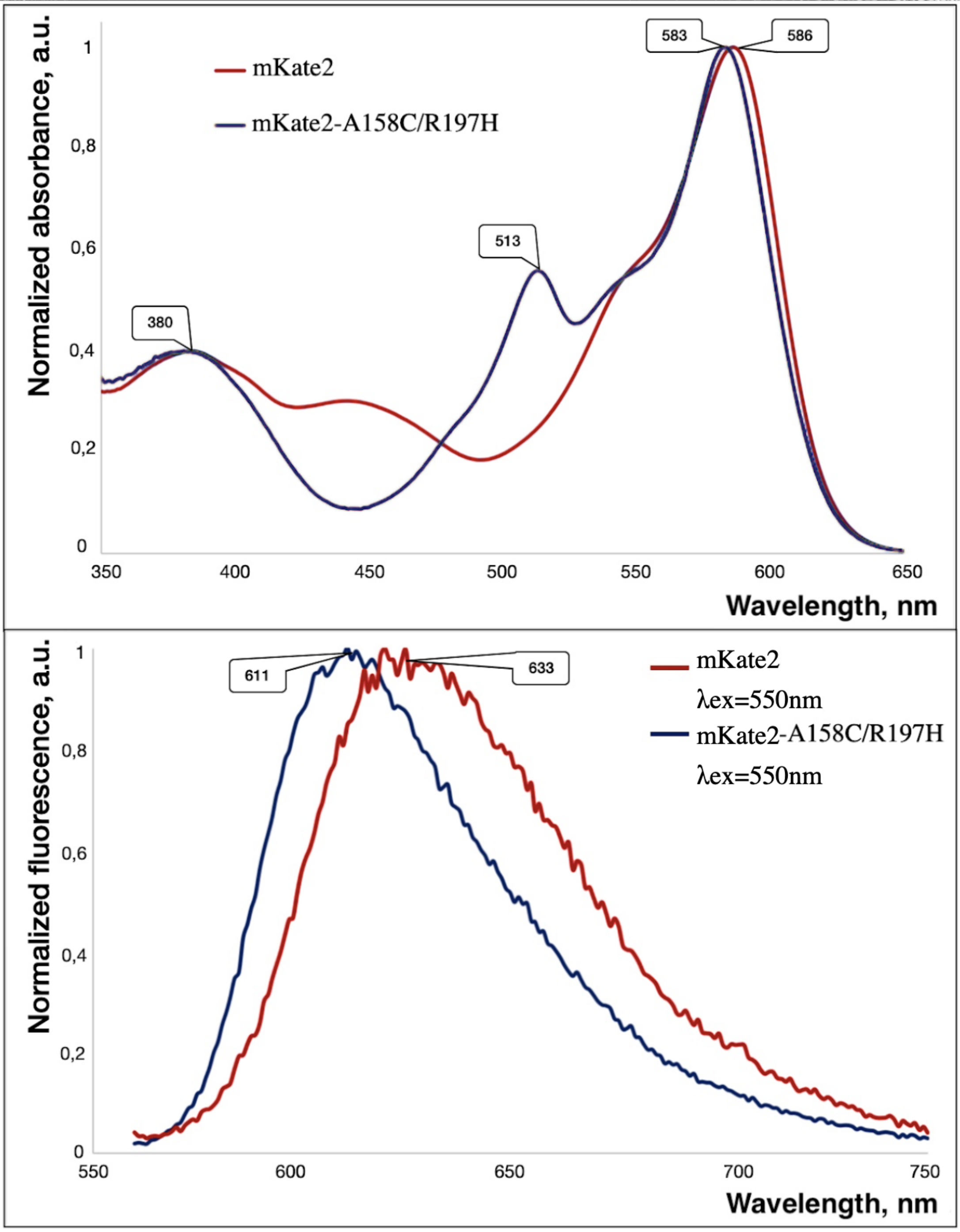


**Supplementary Figure S5.** Absorption (A) and fluorescence emission (B) spectra of mKate2-A158C/R197H compared with those of its parent mKate2. Wavelengths of the major bands' maxima are shown in the bubbles. In the fluorescence graph, excitation wavelengths used for the emission spectra recording are shown in the legend (right upper corner).

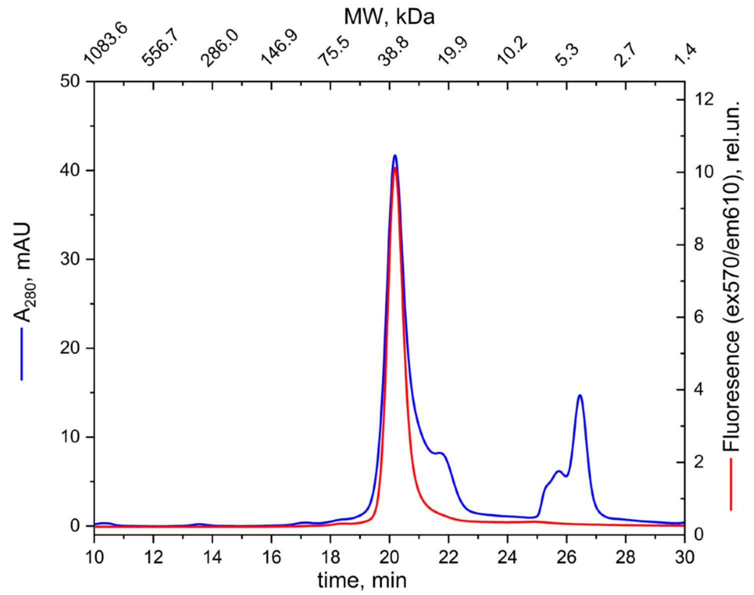


**Supplementary Figure S6**. Gel-filtration chromatography of the purified mKate2-K67R/R197H (Diogenes) sample. Eluate was monitored using in-line absorbance and fluorescence detectors.

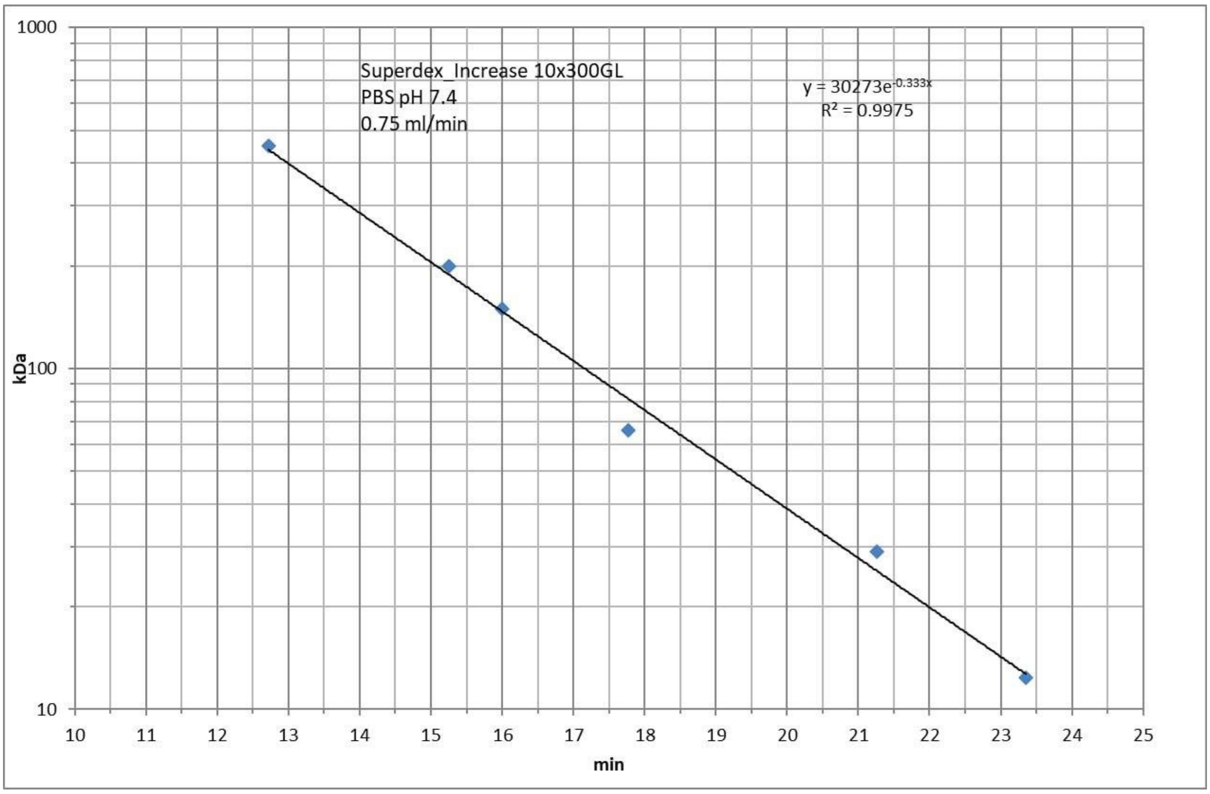


**Supplementary Figure S7.** Calibration plot used to determine molecular weights in the gel-filtration/liquid chromatography setup (see Materials and methods section, main text). Blue squares show the protein standards used for calibration. Column type and equilibration/elution parameters as well as the linear fitting results are described on the graph.

**Supplementary Table S1.** Protein standards used to calibrate a gel-filtration setup. The leftmost column represents protein name, the central – molecular weight, kDa, the rightmost one – elution time while calibrating.

| | | |
|---|---|---|
| CytochromeC horse heart | 12.4 | 23.354 |
| carboanhydrase bovine erytrocytes | 29 | 21.265 |
| BSA | 66 | 17.767 |
| alcohol dehydrogenase yeast | 150 | 15.995 |
| b-amilase sweet potato | 200 | 15.253 |
| ferritin horse spleen | 450 | 12.723 |

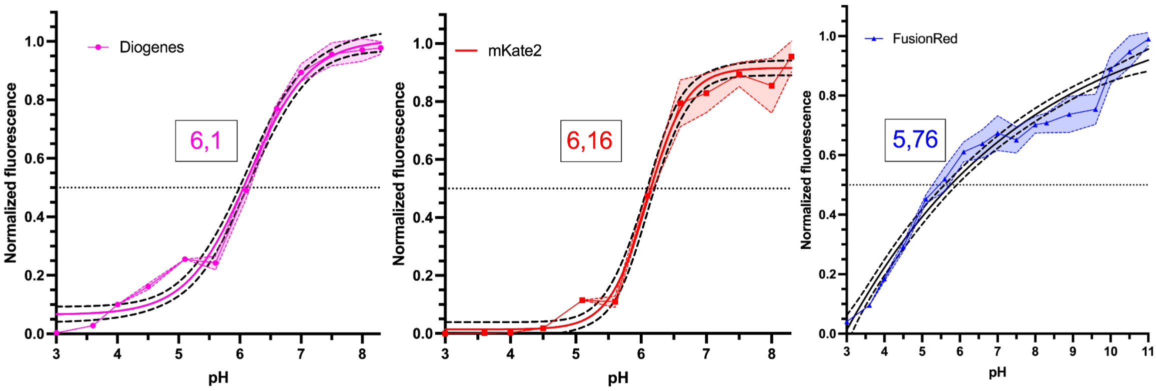


**Supplementary Figure S8.** Sigmoidal fits of the pH-stability data for Diogenes, mKate2 and FusionRed. Colored graphs represent experimental data with standard deviations shown as semi-transparent areas (n=6). Solid black line is the fitting curve with 95% confidence bands shown in black dashed lines. Fitting was performed in the GraphPad Prism10 package (four-parameter logistic curve (4PL) fitting mode was used). Horizontal line stands for half-maximum fluorescence level.

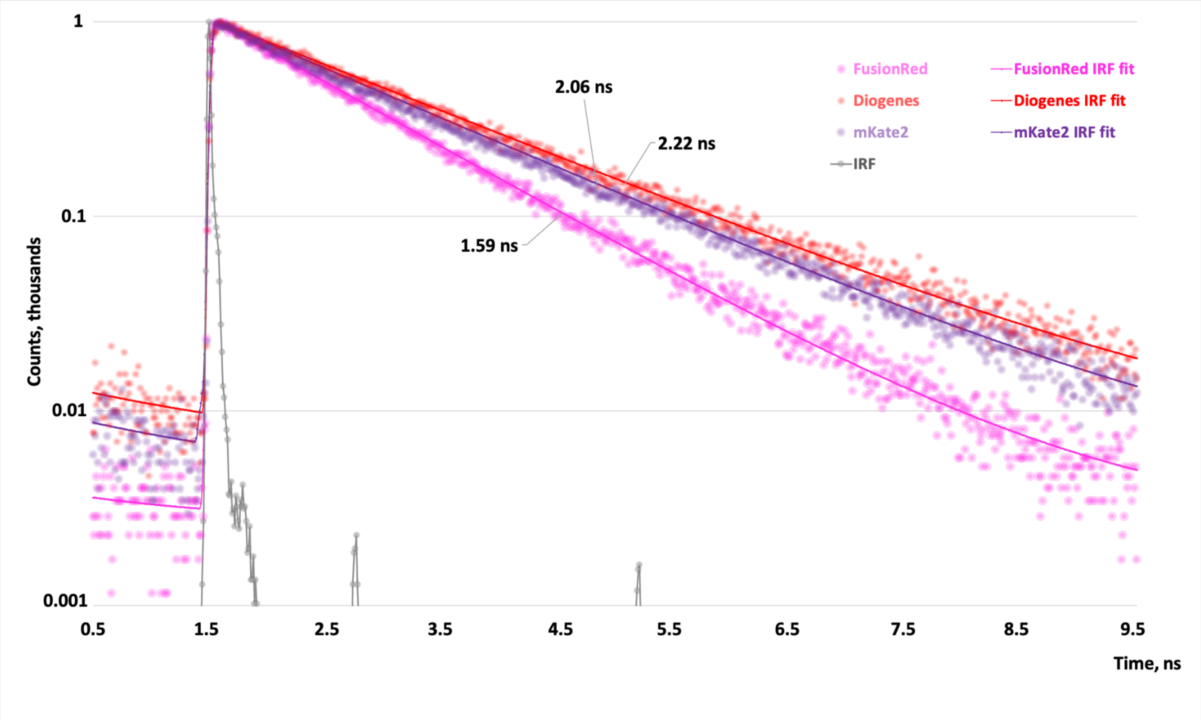


**Supplementary Figure S9.** Fluorescence decay kinetics recorded upon 590 nm single-photon excitation with the femtosecond laser driven at a repetition rate of 80 MHz. The raw-data and exponential fitting curves for FusionRed (biexponential fit), mKate2 and Diogenes (monoexponential fits) are shown. Mean lifetime values are inscribed near the decay curves. Deconvolution with IRF was used for fitting. Measured instrument response (IRF) is shown in grey.

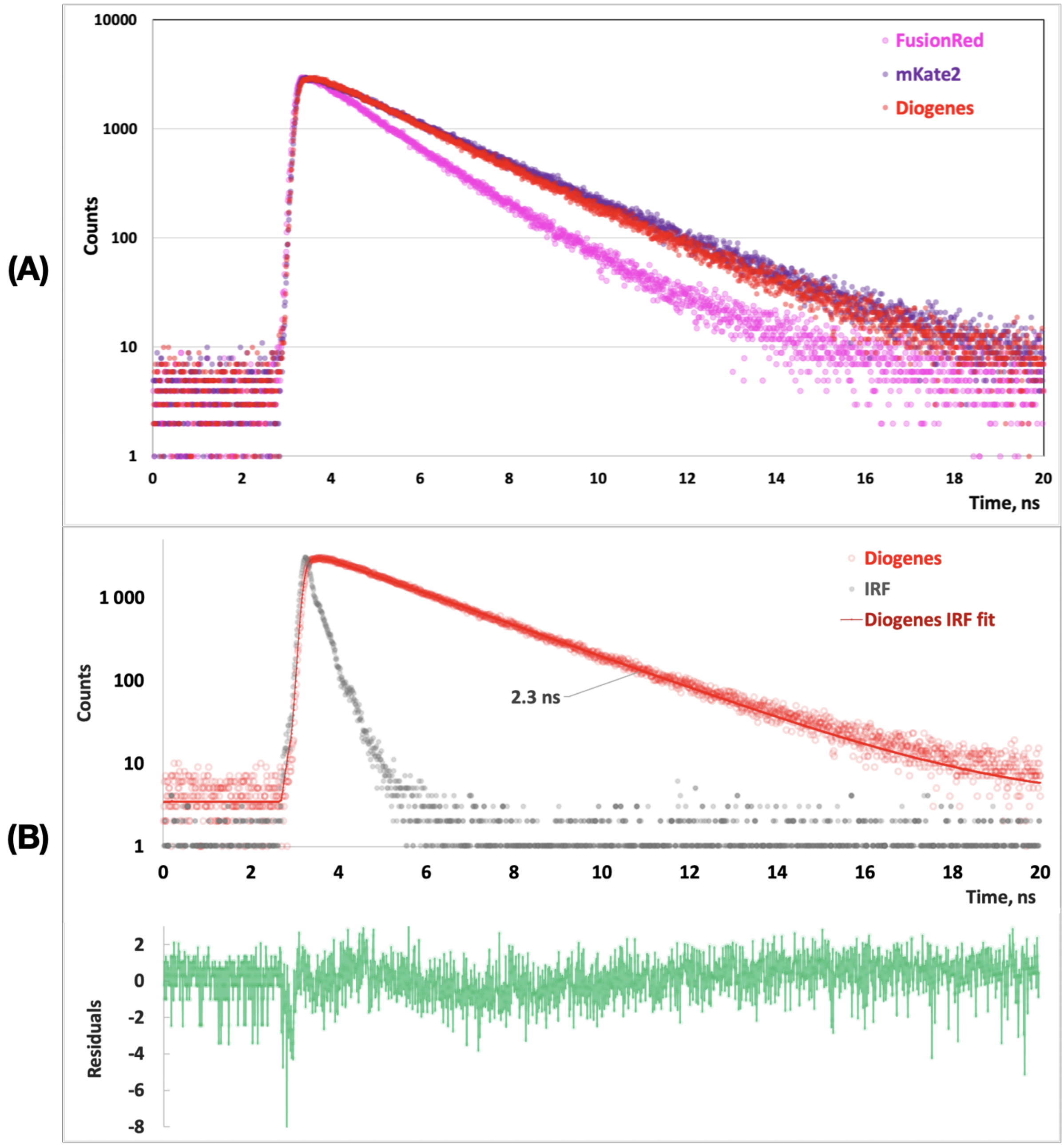


**Supplementary Figure S10.** Fluorescence decay kinetics recorded upon 450 nm single-photon excitation with the picosecond diode laser driven at a repetition rate of 20 MHz. Comparison of the raw-data decay curves for FusionRed, mKate2 and Diogenes (A). Single-component exponential fitting of the Diogenes decay curve (B). Deconvolution with IRF was used for fitting. Measured instrument response (IRF) is shown in grey.

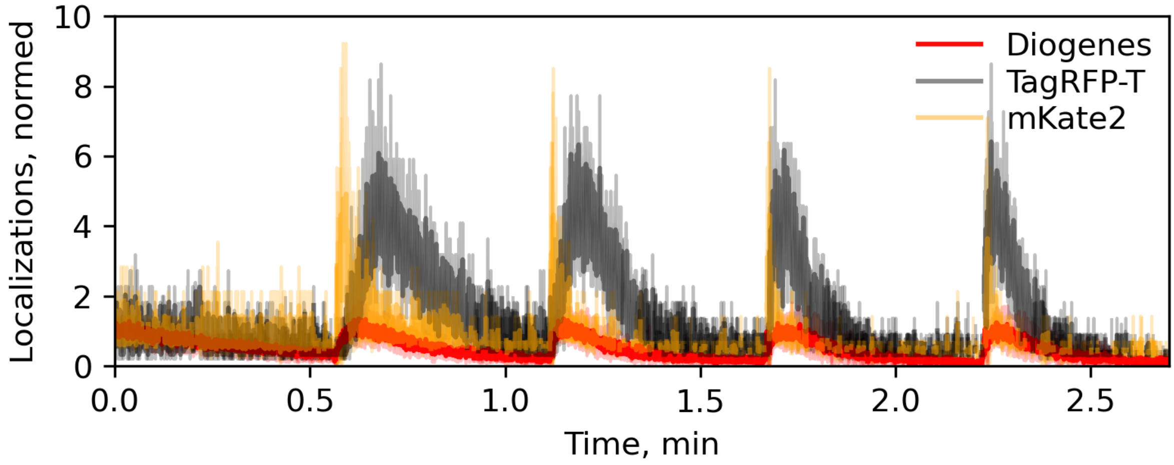


**Supplementary Figure S11.** Effect of UV laser illumination on the localization density of Diogenes, TagRFP-T, and mKate2 as parts of vimentin fusion proteins in live HeLa cells during imaging under the following conditions: 2 kW cm$^{-2}$ 561 nm laser, 16.7 ms frame time, 10,000 frames.
